# Microgravity Remodels Longevity Networks in Astronaut PBMCs: Integrated Findings of Telomere Elongation, DNA Repair Responses, and miRNA Suppression

**DOI:** 10.64898/2025.12.20.695716

**Authors:** Ebru Cam, Ozge Demir, Busra Tekirdagli, Dilara Bulut, Elanur Guzenge, Seden Nadire Harputoglu Efendi, Batuhan Yolver, Kursat Kolay, Ibrahim Halil Kavakli, Fathi Karouia, Beyza Aydin, Cihan Tastan

## Abstract

Microgravity provides a unique environment for elucidating the fundamental mechanisms of human aging. In the Microgravity Associated Genetics (MESSAGE) Science Mission—Turkiye’s first human space biology experiment—we performed an integrative analysis of telomere dynamics, transcriptomic remodeling, and microRNA regulation in peripheral blood mononuclear cells (PBMCs) collected before launch (L-7day), after suborbital ascent (L+3hrs), and during days 4–10 aboard the International Space Station (ISS). Spaceflight induced a striking early elongation of telomeres, accompanied by transcriptional activation of DNA repair, oxidative stress mitigation, mitochondrial homeostasis, and immune regulatory pathways. Concurrently, microgravity triggered robust suppression of longevity-associated microRNAs, including members of the miR-17-92, miR29, and miR34 families, suggesting coordinated epigenetic reprogramming of genome stability and stress responses. Notably, the adaptor protein gene AP2A1, recently implicated in cellular rejuvenation and mechanotransductive aging processes, emerged as a consistently microgravity-responsive hub, linking cytoskeletal signaling to telomere maintenance and DNA repair networks. Together, these findings reveal that short-duration spaceflight initiates a multi-layered molecular longevity program in human immune cells, characterized by telomere extension, stabilization of genome maintenance pathways, and suppression of aging-associated miRNA regulators. This systems-level view provides foundational insight into how human biology adapts to short-term microgravity exposure and identifies AP2A1-centered networks as promising targets for enhancing astronaut health and performance and ultimately understanding terrestrial aging.

## 1. Introduction

Aging is a multifactorial physiological process characterized by the progressive deterioration of cellular structure and function over time. It is associated with key molecular and cellular hallmarks such as telomere shortening, oxidative stress, mitochondrial dysfunction, chronic low-grade inflammation, and genomic instability ^1^. Advances in aging biology have revealed that these processes are not only passive consequences of time but are driven by defined genetic and molecular mechanisms. In this context, longevity-associated genes, including SIRT1, TP53, IL6, and SOD2, have been intensively studied due to their roles in regulating essential cellular pathways such as DNA repair, apoptosis, inflammatory responses, mitochondrial biogenesis, and antioxidant defense ^2–4^.

One of the most intriguing frontiers in aging research is the investigation of how extreme environmental conditions—such as spaceflight—modulate these molecular processes. The microgravity environment encountered during space missions represents a unique physiological stressor, which can alter cytoskeletal organization, impair mitochondrial function, and reprogram immune responses ^5,6^. These effects collectively accelerate or restructure key biological pathways associated with aging ^7^. Evidence from large-scale omics studies, particularly NASA’s Twins Study, has demonstrated that long-term space travel induces significant changes in telomere dynamics, epigenetic regulation, gene expression, and immune system programming, highlighting the bidirectional relationship between the space environment and cellular aging mechanisms. Among the most striking findings of the Twins Study was the transient elongation of telomeres in the astronaut exposed to spaceflight, a paradoxical observation that has challenged traditional views of telomere biology and raised questions regarding the role of microgravity and space-induced stress in genomic stability ^8^. While these observations represent a significant milestone, existing space biology studies have largely focused on individuals of Western origin, and there is a paucity of data from subjects with different ethnic or genetic backgrounds. This constitutes a critical gap in the literature, particularly as space programs expand globally with the emergence of private astronauts.

Recent civilian spaceflight missions have further highlighted the rapid and systemic biological responses elicited by short-term exposure to microgravity. The Inspiration4 mission, the first all-civilian orbital flight, provided a unique opportunity to investigate molecular adaptations in non-professional astronauts under conditions of brief spaceflight ^9^. Multi-omics analyses from Inspiration4 revealed coordinated alterations in immune regulation, mitochondrial function, oxidative stress pathways, and epigenetic signatures, many of which overlapped with changes previously observed in professional astronaut cohorts. Notably, transient modulation of DNA damage response pathways, inflammatory signaling, and regulatory non-coding RNAs was reported, supporting the concept that early spaceflight induces a conserved, adaptive molecular program rather than cumulative damage alone. These findings underscore that even short-duration missions are sufficient to trigger longevity- and stress-related pathways, emphasizing the importance of defining early regulatory mechanisms—including telomere dynamics, DNA repair activation, and miRNA-mediated control—that may govern long-term health outcomes during and after spaceflight. Importantly, the Inspiration4 crew included individuals from diverse demographic backgrounds, including an African American woman, thereby extending spaceflight-associated molecular observations beyond traditionally studied astronaut populations and highlighting the relevance of these findings across broader human diversity.

Microgravity represents a profound environmental stressor that can reshape gene regulatory networks, including post-transcriptional mechanisms governed by microRNAs (miRNAs). As key regulators of messenger RNA (mRNA) stability, translation, and cellular phenotype, miRNAs orchestrate essential pathways related to stress adaptation, immune function, mitochondrial homeostasis, and aging ^10^. Spaceflight studies have demonstrated that microgravity alters the expression of numerous miRNAs involved in oxidative stress responses, cytoskeletal organization, and DNA damage repair, suggesting that miRNA-mediated regulation constitutes a central component of the cellular adaptation process in space ^11^. For instance, NASA’s Twins Study identified significant spaceflight-associated changes in several circulating miRNAs involved in telomere maintenance, inflammation, and mitochondrial function, positioning miRNAs as dynamic biomarkers of physiological stress in spaceflight conditions ^8^. Experimental microgravity models have further shown dysregulation in miRNAs controlling apoptosis, autophagy, and immune signaling—such as miR-21, miR-29, and the miR-17-92 cluster—highlighting their dual roles as both markers and modulators of aging pathways ^12–14^. Given their sensitivity to environmental perturbations and their broad regulatory capacity, miRNA transcript dynamics offer a powerful lens through which to understand how microgravity reshapes cellular aging, immune stability, and stress resilience during spaceflight.

In response to this gap, the Microgravity Associated Genetics (MESSAGE) project was initiated under the leadership of the Turkish Space Agency (TUA), as part of Turkiye’s first human spaceflight mission. Despite growing evidence that microgravity profoundly influences genomic, epigenomic, and transcriptional landscapes, the regulatory contribution of miRNAs and longevity-associated genes to astronaut physiology remains insufficiently characterized. Most existing spaceflight studies have focused on limited gene panels or short-term responses, leaving critical gaps in our understanding of how integrated molecular networks—spanning telomere dynamics, oxidative stress pathways, immune modulation, and miRNA-mediated regulation—coordinate cellular adaptation to gravitational stress ^8,9^. Moreover, data derived from non-Western populations are virtually absent, preventing comprehensive cross-population comparisons of space-induced aging trajectories. In this context, Turkiye’s first human space biology initiative, the MESSAGE Science Mission, presents a unique opportunity to investigate multi-layered aging mechanisms through the use of transcriptomic, telomeric, and miRNA profiling of astronaut-derived immune cells across both suborbital and orbital microgravity exposures. By integrating these datasets, the present study aims to delineate the molecular signatures of microgravity-induced aging and identify both canonical and novel regulators—including the AP2A1 gene family and key aging-related miRNAs—capable of shaping cellular longevity pathways in space. Through this approach, we seek to establish one of the first comprehensive molecular frameworks describing how microgravity reprograms human aging biology and to contribute valuable insights for safeguarding astronaut health during future long-duration missions.

## 2. Materials and Methods

### Ethical Approval and Consent to Participate

The Microgravity Associated Genetics (MESSAGE) Science Mission proposal (Study eIRB Number: STUDY00000620) was reviewed and approved by the National Aeronautics and Space Administration (NASA) Institutional Review Board (IRB) in accordance with ethical standards and the requirements of the Code of Federal Regulations for the Protection of Human Subjects (NASA 14CFR1230 and FDA 21CFR50 and 56, if applicable) (Approval Date: September 21, 2023). The MESSAGE Science Mission proposal (No: 61351342/020-275) was also approved by the Ethics Commission of Üsküdar University (Approval Date: July 03, 2023) ^15^.

### Sample Collection

Blood samples were obtained from three astronauts who participated in an orbital spaceflight and one astronaut in a suborbital flight, as part of the Axiom-3 and Galactic 07 missions, respectively. Samples were collected at three different time points: before launch (L-7 day), in ISS (L+4day, L+7day, and L+10day) and post-suborbital flight (L+3hr) the respective space missions. Venous blood collection was performed using heparinized tubes, and all samples were immediately transported to the laboratory at 4 °C. Additionally, blood samples from healthy volunteers matched for age and gender were collected and processed in parallel as a control group, using the same protocol. The methodology for the isolation of peripheral blood mononuclear cells (PBMCs) in this study was adapted from a previously validated protocol employed during the MESSAGE Science Mission aboard the Axiom-3 flight on the International Space Station ^15^.

### Transcriptome Analysis

Single-stranded complementary DNA (cDNA) was synthesized from total RNA using the Applied Biosystems™ High-Capacity cDNA Reverse Transcription Kit (Thermo Fisher Scientific), following the manufacturer’s protocol to ensure high-quality reverse transcription. The kit components were stored at –20 °C, and all reagents were thawed on ice immediately before use to preserve enzymatic activity. For each reaction, a maximum of 2 μg of RNA was used. The RNA samples were prepared in RNase-free tubes using nuclease-free water or PCR-grade buffer. A 2X reverse transcription master mix was prepared on ice, containing 10X RT Buffer, 25X dNTP Mix (100 mM), 10X Random Primers, MultiScribe™ Reverse Transcriptase, and nuclease-free water. If RNase contamination was suspected, an RNase inhibitor (1 U/μL final concentration) was added to the mix. Equal volumes (10 μL) of the 2X master mix and RNA sample were combined to reach a final reaction volume of 20 μL. The mixture was gently homogenized by pipetting, briefly centrifuged to remove air bubbles, and kept on ice until thermal cycling. Reverse transcription was carried out under the following thermal conditions: 25 °C for 10 minutes, 37 °C for 120 minutes, and 85 °C for 5 minutes to inactivate the enzyme. The resulting cDNA was stored at –20 °C until used in downstream analyses. RNA-Seq analyses were performed on high-quality total RNA samples by SZA OMICS. RNA isolation was conducted using the QIAsymphony (Qiagen, Germany) with the PAXgene Blood RNA Kit (Qiagen; Cat. No. 762431). Library preparation was automated using the Hamilton NGS STAR system with the Illumina Stranded Total RNA Prep and Ribo-Zero Plus Kit (Illumina; Cat. No. 20040529). Paired-end sequencing was performed on the Illumina NovaSeq 6000 platform (2×100 bp) with an average read depth of ∼20–25 million reads per sample. Quality control was performed with FASTQC v0.11.9, and low-quality bases and adapter sequences were removed with TrimGalore v0.6.7. Cleaned reads were aligned to the human reference genome GRCh38 using HISAT2 v2.2.1, and read counts were obtained with HTSeq v0.13.5. Differential gene expression analyses were conducted with DESeq2 v1.34.0 in the RStudio (R v4.2.2) environment.

### Telomere Length Measurement

Telomere length analysis was performed by quantitative real-time PCR (qPCR) using Absolute Human Telomere Length Quantification qPCR Assay Kit (AHTLQ, Cat#8918) developed by ScienCell Research Laboratories (ScienCell, 2023). For this purpose, primers specific for telomere sequences and single-copy reference gene (36B4) primers were used. The reaction mixture was made up to 25 µL with 12.5 µL SYBR Green Master Mix, 1 µL primer mix (10 µM each), 2 µL cDNA, and RNase-free water. PCR conditions: Initial denaturation at 95°C for 10 minutes, followed by 40 cycles of 95°C for 15 seconds and 60°C for 1 minute. Each sample was run in triplicate. Relative telomere length was determined by calculating the T/S ratio (telomere/single copy gene) from Cq (threshold cycle) values. Measurements were performed on the Roche LightCycler® 480, and analyses were calculated automatically with the instrument’s software.

### Statistical Analysis

The Kruskal-Wallis test was applied to data that did not show normal distribution ^16^. The analysis was performed on gene expression levels and telomere length data obtained at before launch (L-7 day), in ISS (L+4day, L+7day, and L+10day) and post-suborbital flight (L+3hr) time points. Statistical significance was accepted as p<0.05. GraphPad Prism 9 software was used for data analysis and graphical presentation. In addition, heatmap visualizations were generated for significant genes, and change profiles were revealed according to time points. Telomere analysis results were also evaluated with group means and standard deviations.

## 3. Results

### Experimental Overview of the MESSAGE Space Biology Mission

To evaluate the effects of microgravity on gene expression associated with immunity and aging, a comprehensive transcriptomic analysis was conducted within the framework of the MESSAGE (Microgravity Associated Genetics Research) project, Turkiye’s first human space biology investigation. In this study, peripheral venous blood samples were obtained from four astronauts—three participating in an orbital mission (Axiom-3) and one in a suborbital flight (Galactic-07)—at multiple time points encompassing pre-flight baseline, in-orbit exposure (ISS days 4, 7, and 10), and acute post-suborbital recovery, thereby capturing distinct gravitational conditions. Samples obtained seven days before launch ( L–7 Days) were preserved at –80 °C and served as baseline controls. Suborbital microgravity was evaluated during the Galactic-07 mission, where samples were collected immediately before and after the flight at approximately 100 km altitude, representing acute exposure conditions. Further sampling occurred on board the International Space Station (ISS) on mission Days 4, 7, and 10, with immediate freezing via the MELFI (Minus Eighty-Degree Laboratory Freezer for ISS) system to ensure RNA integrity. All samples were consistently stored at –80 °C and subsequently processed for high-throughput RNA sequencing (RNA-Seq) to identify transcriptomic shifts under variable gravitational conditions. The full temporal and spatial organization of the experimental sampling strategy is schematically represented in **Figure 1**, which outlines the trajectory from Earth-based controls to suborbital and orbital sampling phases.

**Figure 1:**
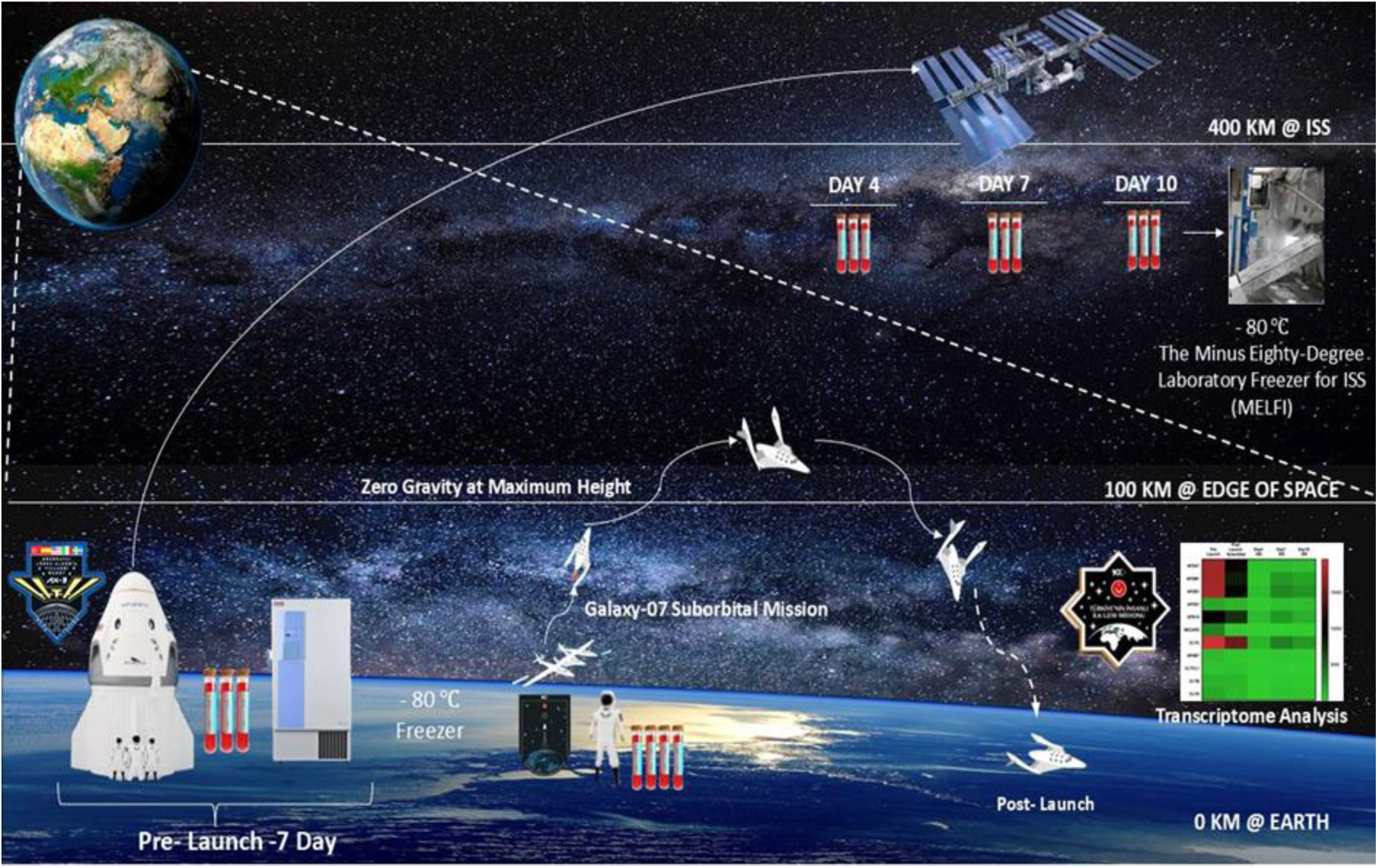
Conceptual framework of the MESSAGE Science Mission. The schematic summarizes blood sample collection and analysis across spaceflight phases. Samples were collected pre-launch on Earth, post-suborbital exposure (∼100 km), and during orbital flight aboard the ISS on days 4, 7, and 10 (∼400 km). All samples were cryopreserved at −80 °C (in-flight samples stored in ISS MELFI) and later analyzed by transcriptomic profiling. This design captures time-dependent transcriptional responses to suborbital and orbital microgravity.

### Global Transcriptomic Shifts in Response to Spaceflight

Transcriptomic profiling of peripheral blood mononuclear cells (PBMCs) collected at multiple time points—Pre-Launch, Post-Launch (Suborbital), and Days 4, 7, and 10 aboard the International Space Station (ISS)—revealed significant transcriptional fluctuations. Data comparisons across astronauts and healthy donors highlighted distinct gene expression trajectories induced by microgravity and suborbital conditions. Differential Expression of Longevity-Associated Genes: A panel of 49 genes, including well-characterized longevity regulators and the AP2A1 gene family, was analyzed. Expression profiles were compared using both statistical (Kruskal-Wallis test) and visual (heatmap) methods to assess responses to altered gravity.

### Differential Regulation of the AP2A1 Gene Family and Endocytic Machinery Under Microgravity

To investigate whether microgravity influences the molecular components of the clathrin-mediated endocytic pathway, expression profiles of the AP2A1 gene family and its associated adaptor and clathrin subunits were examined across all mission phases (**Fig. 2**). A STRING-based protein–protein interaction analysis revealed a highly interconnected network encompassing AP2 complex subunits (AP2A1, AP2M1, AP2B1, AP2S1), clathrin light and heavy chains (CLTA, CLTB, CLTC), and accessory proteins (NECAP1, NECAP2, EPS15, AP4B1), underscoring their coordinated involvement in vesicle formation and intracellular trafficking ^9,17^. Across nearly all genes analyzed, a consistent and marked reduction in transcript abundance was observed following exposure to both suborbital and orbital microgravity. AP2A1 expression decreased significantly from baseline to post-flight conditions (p = 0.0082), with similar suppression evident for AP2M1 (p = 0.0056), AP2B1 (p = 0.0047), and AP2S1 (p = 0.0366). Clathrin-related genes also demonstrated pronounced downregulation, including CLTC (p = 0.0018), CLTCL1 (p = 0.0010), and CLTB (p = 0.0047). Accessory proteins NECAP2 (p = 0.0308), AP4B1 (p = 0.0018), and EPS15 (p = 0.0023) followed the same decreasing trajectory. Notably, transcript levels showed the greatest decline immediately after suborbital exposure and remained suppressed throughout Days 4, 7, and 10 aboard the ISS, with minimal recovery over time. This pattern indicates that both acute and sustained microgravity environments impose significant regulatory pressure on endocytic pathway genes ^9,11,17^. The uniformity of suppression across multiple functional components—including adaptor proteins, clathrin subunits, and regulatory cofactors—suggests a coordinated transcriptional reprogramming of vesicular trafficking machinery in response to gravitational unloading ^11,18^. Together, these findings demonstrate that microgravity robustly downregulates the AP2A1 gene family and its associated endocytic network, supporting the hypothesis that alterations in membrane trafficking and mechanotransduction represent fundamental components of the cellular adaptation to spaceflight.

**Figure 2.**
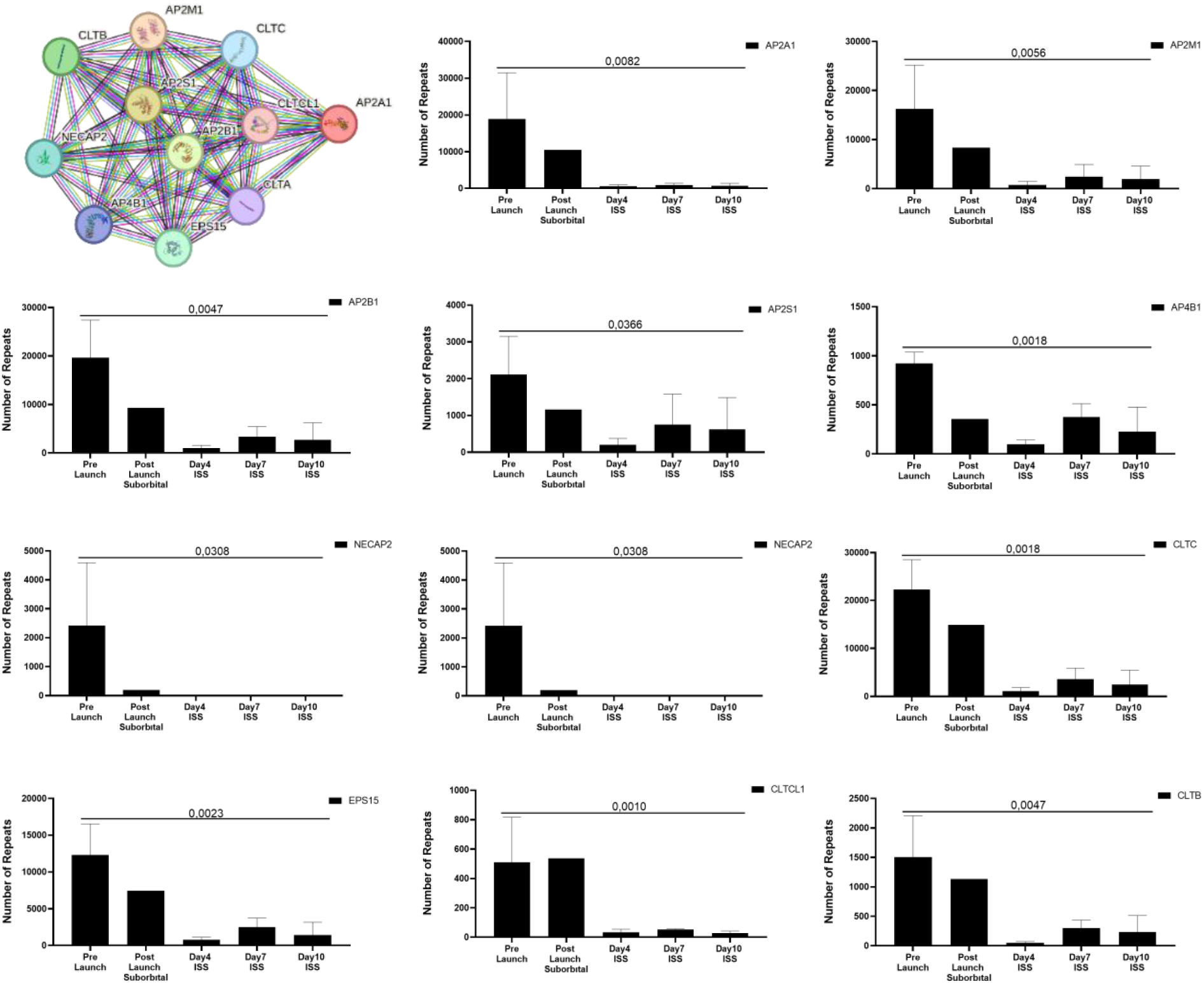
Differential expression and interaction network of the AP2A1 gene family under microgravity and suborbital conditions. This figure illustrates the transcriptional response of the AP2A1 gene family and associated endocytic adaptor proteins across five mission time points: Pre-Launch (baseline), Post-Launch Suborbital, and Days 4, 7, and 10 aboard the ISS. (Left) Protein–protein interaction (PPI) network generated via STRING demonstrates functional connectivity among AP2A1 family members (AP2A1, AP2A2, AP2B1, AP2S1), clathrin components (CLTA, CLTB, CLTC), accessory proteins (NECAP1/2), and EPS15, highlighting their coordinated roles in clathrin-mediated endocytosis. (Right) Bar graphs show normalized RNA-Seq read counts for each gene across mission phases. Significant decreases in expression between Pre-Launch and subsequent microgravity time points were detected for multiple AP2 complex genes (AP2A1, AP2M1, AP2B1, AP2S1), clathrin heavy and light chain genes (CLTC, CLTA, CLTB), and adaptor proteins (AP4B1, NECAP1/2, EPS15), as determined by the Kruskal–Wallis test (p-values indicated above each panel). Error bars represent standard deviation. Collectively, these data demonstrate a broad suppression of endocytic machinery components under microgravity, suggesting a gravity-sensitive regulatory shift in membrane trafficking pathways during spaceflight.

### Heatmap Analysis Reveals Coordinated Downregulation of Endocytic Machinery in Microgravity

Heatmap visualization of AP2 complex members and clathrin-associated genes revealed a robust and synchronized decrease in expression following exposure to microgravity (**Fig. 3**). Baseline (Pre-Launch) samples showed high transcript abundance for AP2A1, AP2M1, AP2B1, CLTC, and related adaptor proteins; however, expression dropped sharply after the suborbital flight and remained consistently low across Days 4, 7, and 10 on the ISS. This pattern was particularly evident for AP2A1, AP2M1, AP2B1, CLTC, and EPS15, indicating that both core adaptor subunits and clathrin heavy chain components are strongly gravity-responsive. Genes such as NECAP2, AP4B1, and clathrin light chain isoforms also exhibited sustained suppression, suggesting that microgravity exerts system-wide effects on vesicle trafficking networks rather than isolated transcriptional shifts ^11^. Together, these results support a coordinated downregulation of endocytosis-related pathways under microgravity, consistent with the disruption of cytoskeletal tension, membrane dynamics, and mechanotransduction cues during spaceflight.

**Fig. 3.**
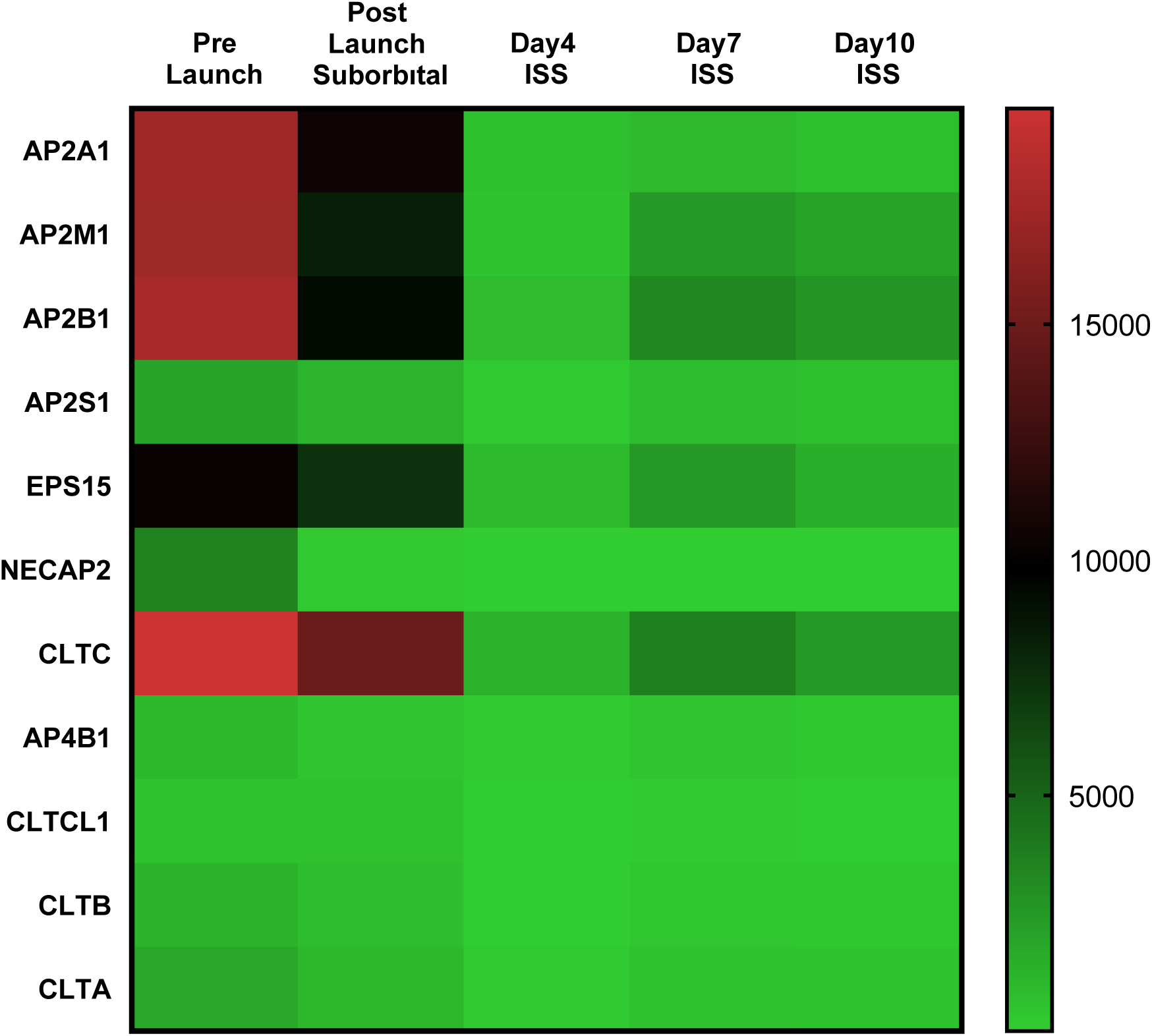
Heatmap of AP2 complex and clathrin-associated gene expression across microgravity exposure conditions. This heatmap illustrates normalized RNA-Seq expression levels of AP2 complex subunits (AP2A1, AP2M1, AP2B1, AP2S1), endocytic accessory proteins (EPS15, NECAP2, AP4B1), and clathrin components (CLTC, CLTCL1, CLTB, CLTA) across five mission phases: Pre-Launch (baseline), Post-Launch Suborbital, and Days 4, 7, and 10 aboard the ISS. Red shading represents higher expression values, while green indicates reduced expression. A pronounced decrease in transcript abundance is observed immediately following suborbital microgravity exposure, with expression remaining suppressed throughout all orbital time points. The consistent downward shift across AP2-family genes, clathrin heavy/light chain isoforms, and associated adaptors highlights a coordinated repression of endocytic machinery under microgravity conditions.

### Microgravity Induces Widespread Downregulation of Longevity-Associated Gene Networks

To characterize how microgravity alters molecular pathways involved in cellular aging, we analyzed expression profiles of 45 longevity-related genes across all mission phases (**Fig. 4**). The data revealed a striking global suppression of transcripts associated with genome maintenance, metabolic regulation, oxidative stress defense, mitochondrial quality control, and inflammatory signaling. Core longevity regulators such as SIRT1, MTOR, WRN, TP53, BRCA2, and GDF11 exhibited large and statistically significant decreases immediately after suborbital exposure (p < 0.05 for all) and remained suppressed throughout ISS Days 4, 7, and 10. The magnitude of downregulation was particularly pronounced for SIRT1 and MTOR, suggesting disruption of nutrient-sensing, DNA repair, and mitochondrial homeostasis pathways under microgravity^1^. Oxidative stress and mitochondrial function genes, including NFE2L2, PINK1, ACY2P, and UCPs, also showed significant reductions post-flight. This pattern is consistent with impaired mitochondrial turnover, reduced antioxidant capacity, and altered ROS-mediated signaling—hallmarks of both aging and spaceflight-induced cellular stress^1,11,19^. Genes associated with DNA damage responses (ATM, DCAF8) and senescence-associated secretory pathways (NLRP3, IL1B, TNFSF10) similarly displayed persistent downregulation across orbital time points. This suggests a dampened inflammatory and DNA repair landscape that may influence both immune reactivity and genomic stability during gravitational unloading^5,11^. Members of the apoptosis and cell survival network, including MCL1, BCL2, FAS, and HMGB1, were significantly reduced, indicating suppression of both pro-survival signaling and programmed cell-death regulation. Notably, mitochondrial apoptotic regulators such as MCL1 showed near-complete silencing by Day 4 of the ISS mission. A distinct pattern emerged for IL15, which was downregulated in early post-flight samples before declining during orbital sampling. IL15 plays a key role in NK-cell and T-cell homeostasis, suggesting a microgravity-triggered compensatory immune mechanism^5,20^. Genes influencing angiogenesis and tissue remodeling, including VEGFA, VEGFB, TGFA, and HGF, were also significantly reduced after launch, implying potential impacts on vascular integrity and regenerative capacity in microgravity. Finally, regulators of mechanotransduction and circadian control—such as CLOCK, GSK3B, and MAPK8—displayed marked suppression, providing evidence that gravitational unloading disrupts cytoskeletal tension signaling and timing pathways intimately linked to aging physiology^11,17^. Overall, this coordinated multi-gene downregulation demonstrates that microgravity acts as a potent environmental modulator of longevity pathways, attenuating transcriptional programs that maintain genomic stability, mitochondrial efficiency, immune competence, and cellular resilience. These findings support a systems-level reorganization of aging biology during spaceflight and align with emerging literature on gravity-dependent regulation of cellular homeostasis^8,11^. Several genes exhibited non-monotonic expression patterns across the ISS time points. Notably, IFNG and BAG3 showed transient upregulation at mission day 7 compared with both day 4 and day 10, indicating a mid-flight modulation of immune and stress-response pathways during sustained microgravity exposure. In contrast, expression levels at day 10 returned toward or below earlier in-flight levels, suggesting that these changes were temporally restricted rather than progressive. Distinct from this pattern, CDKN2A displayed a sustained upregulation during spaceflight relative to pre-launch baseline, exhibiting a divergent trajectory compared with most proliferation- and repair-associated genes. This increase was observed across multiple in-flight time points, indicating selective activation of cell-cycle checkpoint regulation during microgravity exposure. Together, these findings highlight temporal heterogeneity in transcriptional responses to spaceflight, with subsets of immune, stress-adaptive, and cell-cycle regulatory genes exhibiting phase-specific modulation rather than uniform directional change.

**Fig. 4.**
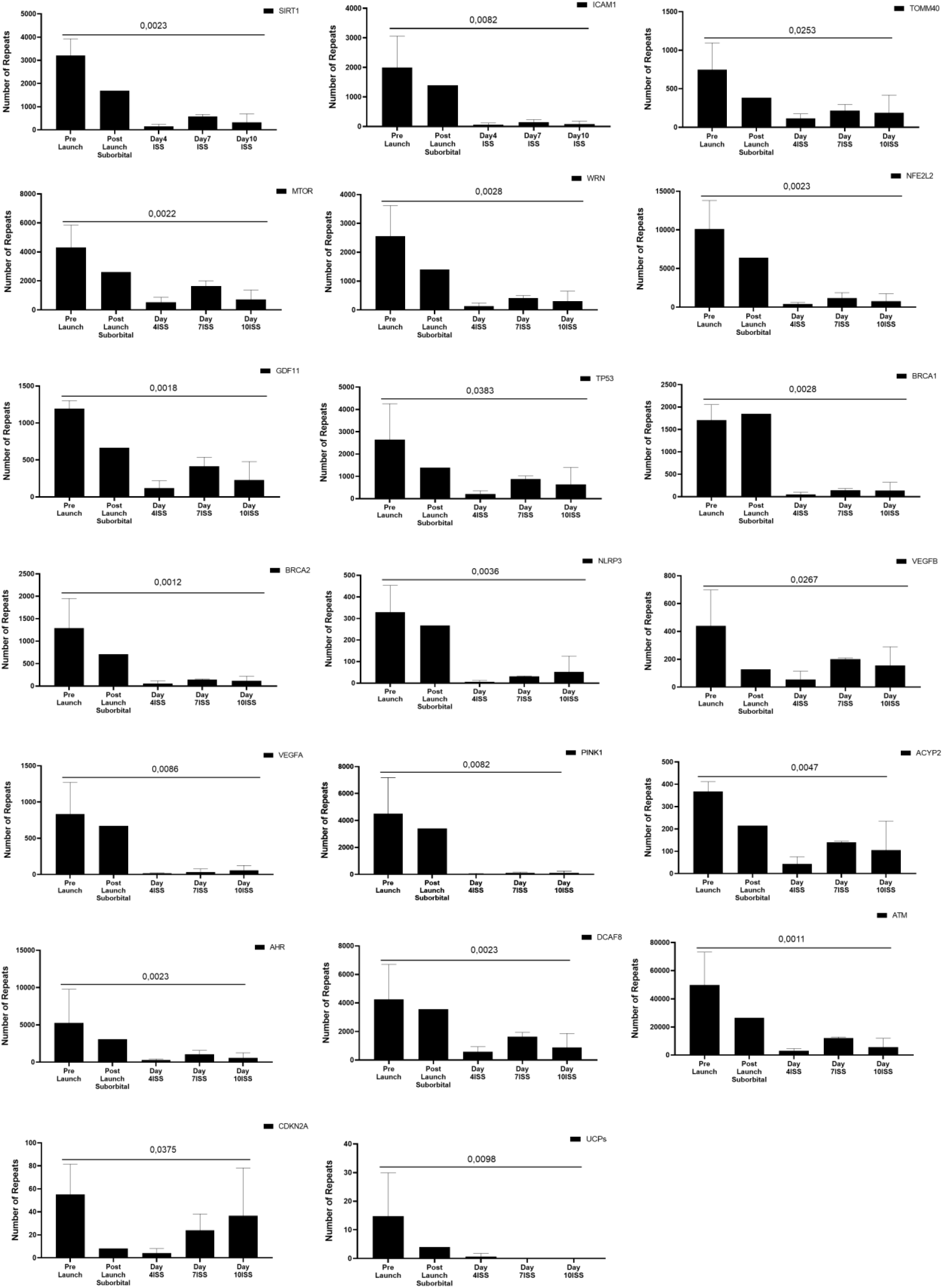
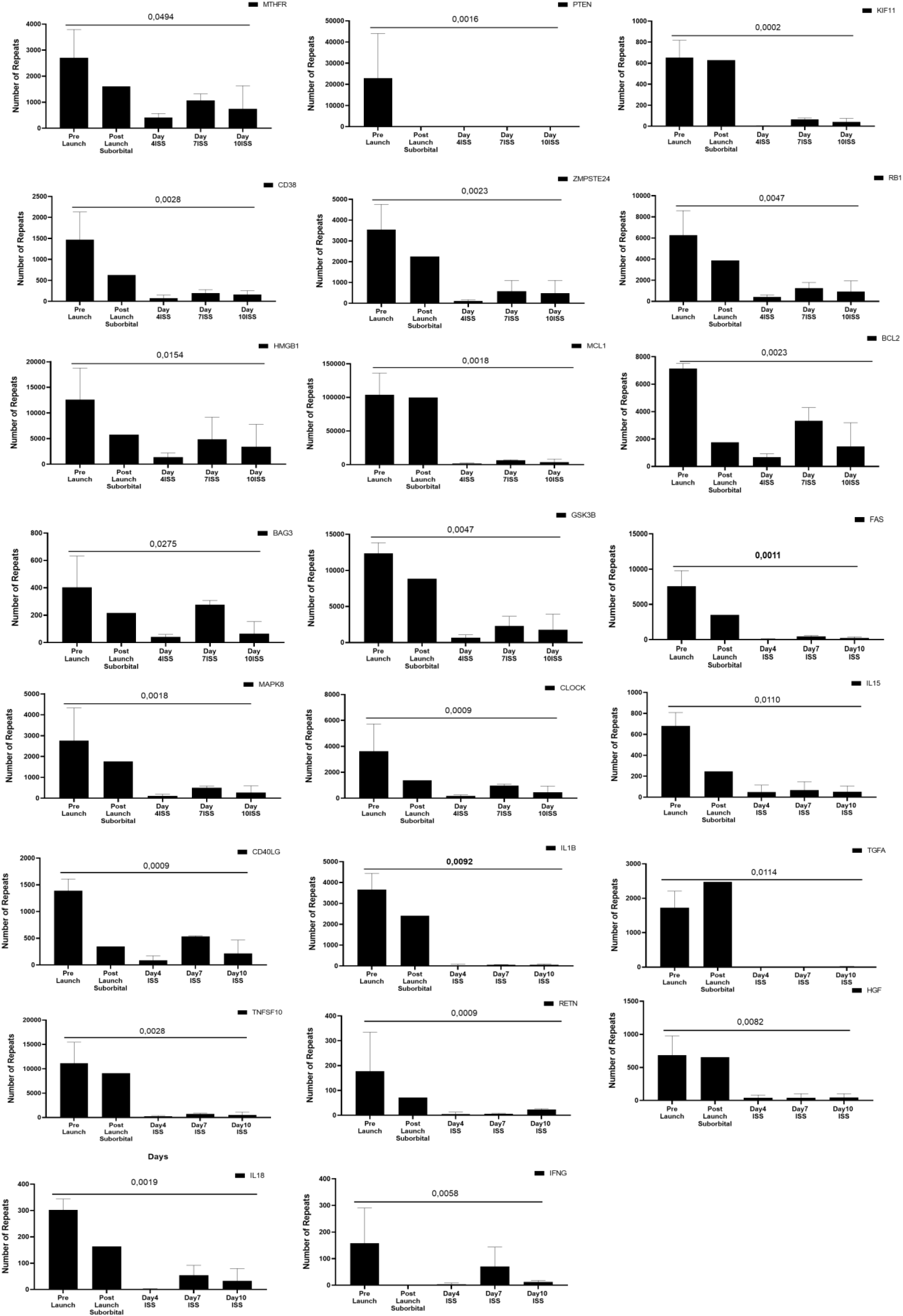
Differential expression of longevity-associated and stress-response genes across suborbital and orbital microgravity exposures. Bar graphs display normalized RNA-Seq read counts for key genes involved in aging, DNA repair, oxidative stress regulation, mitochondrial function, inflammatory signaling, mechanotransduction, and immune homeostasis. Expression levels are shown for five mission phases: Pre-Launch (baseline), Post-Launch Suborbital, and Days 4, 7, and 10 aboard the ISS. For each gene, Kruskal–Wallis p-values are indicated above the plots. The first panel set includes canonical longevity regulators (SIRT1, MTOR, WRN, TP53, GDF11, BRCA1/2), oxidative stress and antioxidant response genes (NFE2L2, PINK1, UCP1), senescence-associated inflammatory mediators (NLRP3, IL1B, TNFSF10, IFNG), angiogenesis-related genes (VEGFA, VEGFB, TGFA, HGF), mitochondrial maintenance genes (ACY2P, CLOCK), and DNA damage response genes (ATM, DCAF8). The second panel set includes additional genes implicated in stress adaptation and aging biology, such as PTEN, MTHFR, KLF11, CD38, HMGB1, MCL1, BCL2, BAG3, MAP3K, CD40LG, ZMPSTE24, GSK3B, and IL15. Error bars represent standard deviations. Across the majority of genes, spaceflight exposure—beginning with suborbital microgravity and persisting through ISS sampling—results in substantial downregulation relative to baseline, with only a limited subset (e.g., CDKN2A) exhibiting increased expression. These findings indicate that microgravity induces a coordinated suppression of multiple molecular pathways central to aging, genome stability, mitochondrial function, and immune regulation (p <0.05, Kruskal-Wallis).

Heatmap analysis revealed the downregulation of longevity-associated genes following exposure to microgravity (**Fig. 5**). Baseline samples showed moderate to high expression across core aging regulators—including SIRT1, MTOR, WRN, GDF11, TP53, and NFE2L2—but these transcripts dropped significantly after suborbital flight and remained uniformly suppressed during ISS Days 4, 7, and 10. This pattern indicates rapid and persistent inhibition of pathways governing genomic maintenance, oxidative stress resistance, nutrient signaling, and mitochondrial quality control^1,11^. Similarly, genes involved in DNA damage responses (ATM, BRCA1, BRCA2, DCAF8) and mitochondrial homeostasis (PINK1, UCP1, TOMM40) showed widespread reductions, consistent with spaceflight-induced alterations in ROS handling, mitophagy signaling, and genome surveillance^8,19^. Notably, MCL1 and HMGB1, both associated with early stress response and apoptosis regulation, exhibited transient post-launch elevation before declining at later ISS time points, suggesting an acute cellular reaction to launch-associated mechanical and metabolic stress^5,11^. Inflammatory and immune-modulatory genes—including IL1B, TNFSF10, IFNG, and ICAM1—also demonstrated significant downregulation, reflecting reduced inflammatory tone and potential immune suppression under microgravity. Downregulation of mechanotransduction-linked genes (GSK3B, CLOCK, MAP3K) further aligns with the disruption of cytoskeletal tension and gravity-dependent signaling cues^11,17^. Altogether, the heatmap demonstrates that microgravity triggers a widespread transcriptional collapse across diverse longevity, mitochondrial, DNA repair, and immune pathways, supporting a model in which gravitational unloading profoundly reshapes molecular programs central to cellular aging and stress adaptation.

**Fig. 5.**
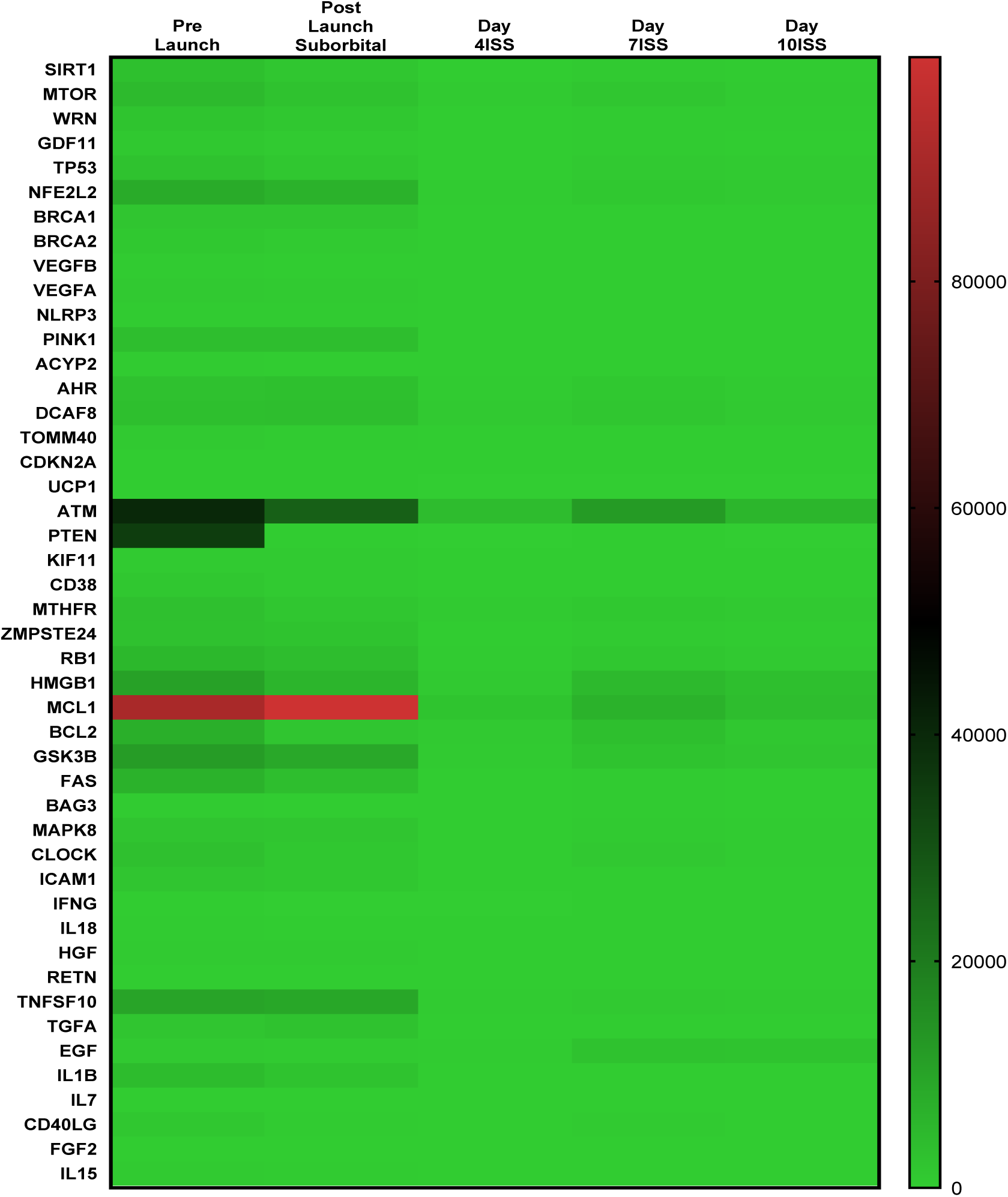
Heatmap of longevity-related, stress-response, and immune regulatory gene expression across suborbital and orbital microgravity conditions. This heatmap displays normalized RNA-Seq expression values for 46 genes involved in cellular aging, genome stability, mitochondrial function, oxidative stress defense, inflammatory signaling, and mechanotransduction across five mission phases: Pre-Launch (baseline), Post-Launch Suborbital, and ISS Days 4, 7, and 10. Red shading represents higher expression, green indicates lower expression, and intermediate shades denote moderate expression levels. A broad transition from moderate baseline expression to markedly reduced transcript abundance is observed following suborbital microgravity exposure, with sustained suppression during all subsequent ISS sampling days. While most longevity and DNA repair genes (e.g., SIRT1, MTOR, WRN, TP53, BRCA1/2, NFE2L2, PINK1) exhibit downregulation, a small subset—including MCL1 and HMGB1—shows transient elevation immediately after launch, suggesting early stress-response activation before overall transcriptional decline. Genes associated with immune activity and cytokine signaling (IL1B, TNFSF10, IFNG, IL15) predominantly show decreased expression, indicating microgravity-dependent dampening of inflammatory pathways. Overall, the heatmap highlights a coordinated, system-wide reprogramming of aging-related molecular pathways in response to microgravity exposure.

### Microgravity Alters Highly Connected Interaction Networks Governing Aging, Genome Stability, Mitochondrial Function, and Immune Signaling

To determine whether transcriptional changes induced by microgravity disrupt broader molecular systems, we examined STRING protein-interaction networks for all significantly affected longevity-related genes (**Fig. 6**). Network analysis revealed that nearly every microgravity-responsive gene occupies a central, highly connected position within its biological module, suggesting that even modest transcript reductions may propagate functional consequences across multiple pathways. Genes implicated in genome maintenance and DNA repair—including ATM, TP53, WRN, BRCA1, BRCA2, CDKN2A, DCAF8, and RB1—formed dense interaction networks enriched for cell-cycle control, checkpoint signaling, and double-strand break repair^1,8^. Because microgravity consistently downregulated these genes, the impact likely extends well beyond individual loci, affecting integrated DNA-repair hubs essential for telomere preservation and genomic stability. Similarly, microgravity-suppressed mitochondrial and metabolic regulators—including TOMM40, PINK1, UCP1, ACYP2, GDF11, and MTHFR—mapped to networks governing mitochondrial import, oxidative phosphorylation, redox defense, and apoptotic priming. These findings align with mitochondrial dysfunction reported in human spaceflight and further support a model in which mechanical unloading disrupts organelle turnover and metabolic longevity pathways^8,19^. Genes regulating nutrient sensing and cellular aging, such as SIRT1, MTOR, MAPK8, and CLOCK, exhibited highly interconnected networks linking circadian regulation, AMPK–mTOR signaling, and metabolic homeostasis. Their coordinated downregulation reflects a shift toward energy-conserving, stress-responsive states characteristic of cells experiencing environmental unloading^11,18^. Microgravity also profoundly affected immune and inflammatory networks, including NLRP3, IL1B, TNFSF10, IL18, IFNG, CD40LG, ICAM1, and RETN, each embedded in dense cytokine, NF-κB, and inflammasome-related networks. Their reduced expression suggests a systematic dampening of innate and adaptive immune activation, consistent with known immune dysregulation in astronauts^5,20^. Finally, apoptosis- and survival-related genes—BCL2, MCL1, FAS, BAG3—showed strong connectivity to caspase signaling and stress-response pathways^1^. Their downregulation points to altered thresholds for cell survival under microgravity conditions.

**Fig. 6.**
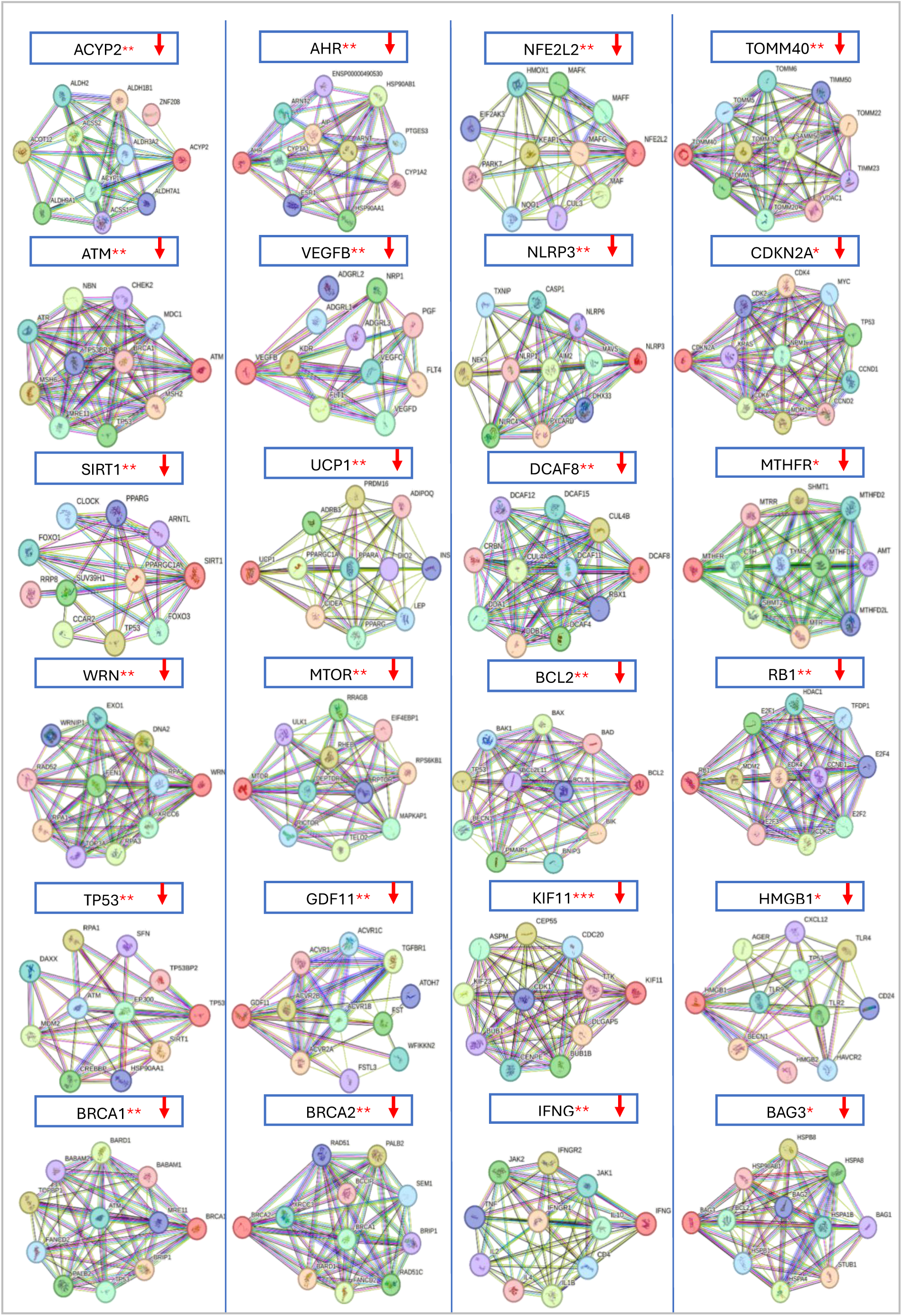
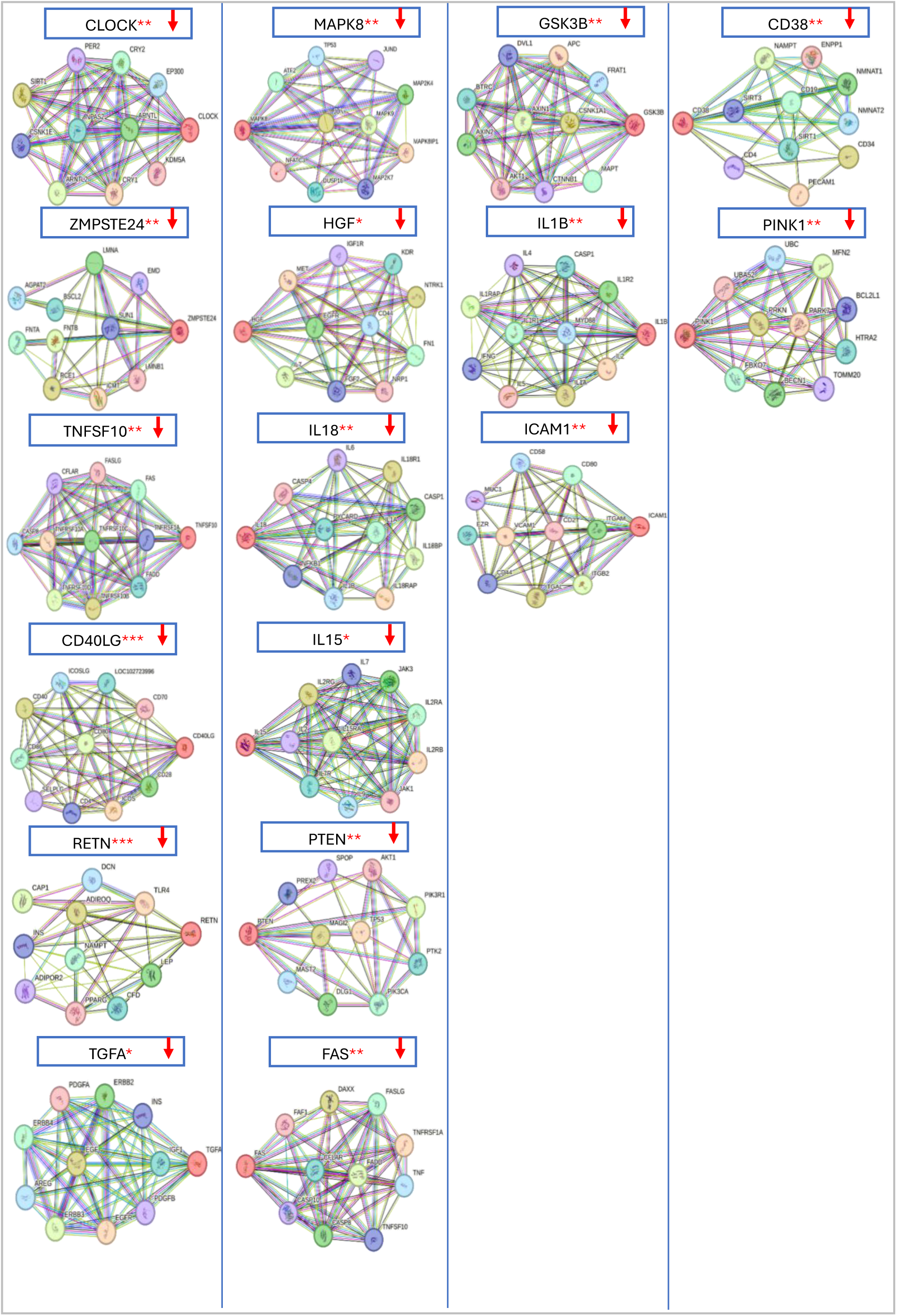
STRING protein–protein interaction (PPI) networks of longevity-, stress-, mitochondrial-, inflammatory-, and DNA-repair–related genes significantly altered by microgravity exposure. Each panel displays a STRING-generated PPI network for an individual gene whose expression was significantly reduced following suborbital and orbital microgravity exposure (indicated by ↓ symbols). Nodes represent proteins; edges depict known or predicted functional associations, including co-expression, experimental evidence, curated pathways, and text-mined interactions. Genes are grouped by biological function, including: Mitochondrial and metabolic regulators: ACYP2, TOMM40, UCP1, PINK1, GDF11, MTHFR; Aging and nutrient-sensing factors: SIRT1, MTOR, CLOCK, MAPK8; DNA damage and genome stability regulators: ATM, WRN, BRCA1, BRCA2, TP53, CDKN2A, DCAF8, RB1; Oxidative stress and detoxification genes: NFE2L2, AHR; Angiogenesis and growth signalling pathways: VEGFA/B, TGFA, HGF; Inflammatory and immune-signaling mediators: NLRP3, IFNG, IL18, IL15, TNFSF10, IL1B, RETN, ICAM1, CD40LG; Apoptosis and cell-survival networks: BCL2, MCL1, FAS, BAG3; Transport and vesicular trafficking components: KIF11, TOMM40, CD38. Red downward arrows indicate significant downregulation compared to pre-launch levels. STRING networks highlight dense connectivity among molecular partners, suggesting that altered expression of each microgravity-responsive gene may disrupt broader biological modules involved in cellular aging.

### Stable Expression of Several Metabolic, Angiogenic, Telomeric, and Inflammatory Genes During Microgravity Exposure

In contrast to the widespread downregulation observed for core longevity, DNA repair, and mitochondrial maintenance genes, a distinct subset of transcripts exhibited no significant change across suborbital or orbital microgravity conditions (**Fig. 7**). Telomerase catalytic subunit TERT, despite its central role in telomere maintenance, showed variable but statistically non-significant fluctuations (p = 0.2135), suggesting that telomerase activity is not acutely transcriptionally regulated in response to microgravity in peripheral immune cells. Similarly, key inflammatory mediators (IL-6, IL-7), metabolic genes (APOE, FTO), angiogenesis-related factors (VEGFC, VEGFD, EGF, FGF2), and vascular tone regulators (ACE) remained transcriptionally stable throughout all sampling points (all p > 0.05). The oxidative stress gene SOD2, extracellular matrix modifiers (TIMP1, SERPINE1), and neuroimmune gene CNR1 also showed no significant expression shifts, indicating that not all stress-response or endocrine pathways respond uniformly to microgravity. Overall, these findings demonstrate that while microgravity exerts profound suppressive effects on longevity, DNA repair, mitochondrial, and immune–inflammatory pathways, a set of metabolic, angiogenic, and cytokine-related genes remains resilient and transcriptionally unchanged. This selective stability may reflect pathway-specific thresholds for gravitational sensitivity or compensatory homeostatic mechanisms that buffer certain biological systems during spaceflight^11,18^.

**Fig. 7.**
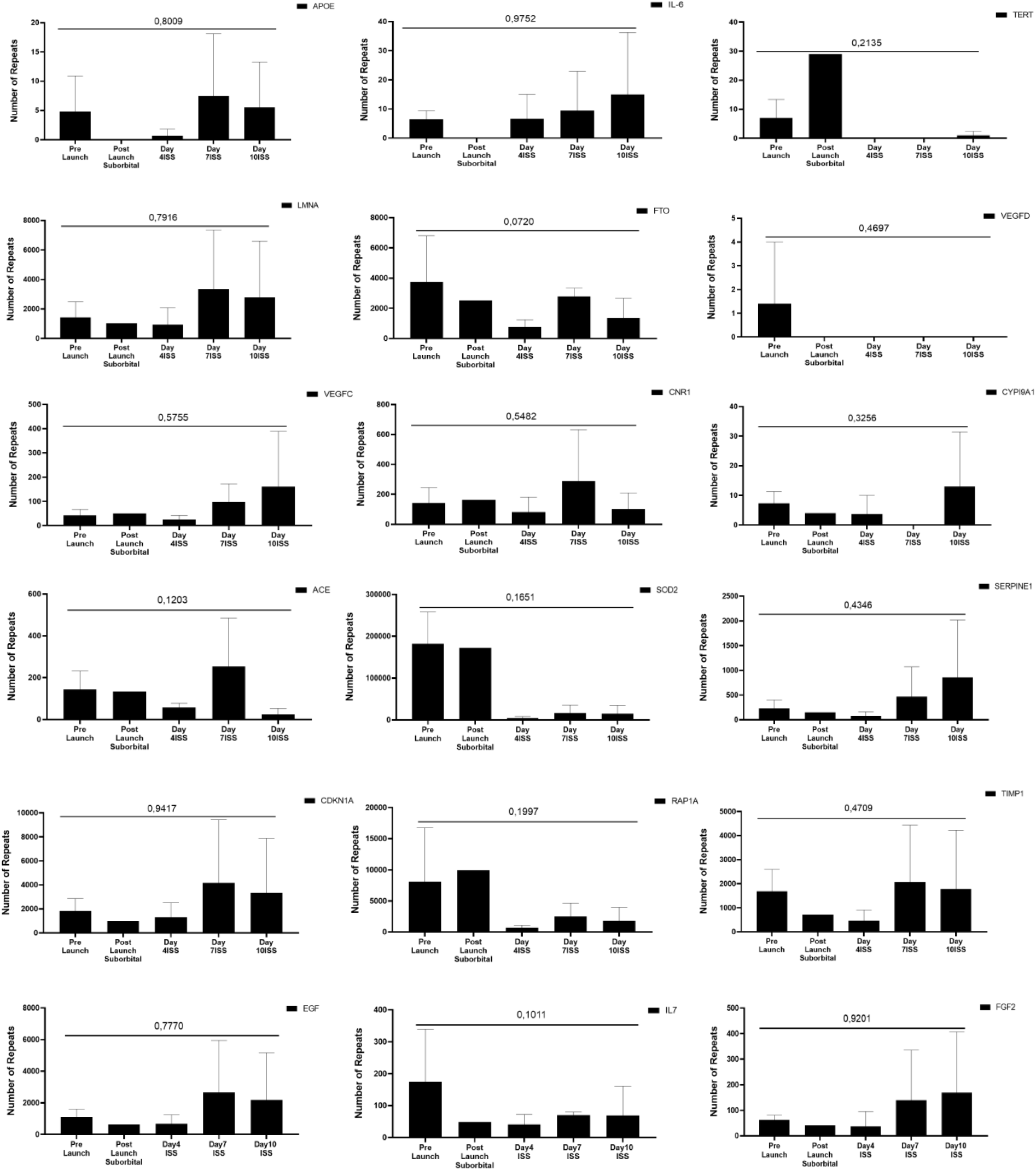
Gene expression profiles of aging-, metabolism-, angiogenesis-, telomere-, and inflammation-related genes that did not show significant changes across microgravity exposure conditions. Bar graphs display normalized RNA-Seq read counts for genes involved in metabolic regulation (APOE, FTO), inflammatory signaling (IL-6, IL-7), telomere biology (TERT), angiogenesis (VEGFC, VEGFD, EGF, FGF2), xenobiotic metabolism (CYP1A1), oxidative stress regulation (SOD2), extracellular matrix remodeling (TIMP1, SERPINE1), neuroendocrine pathways (CNR1), and systemic blood pressure or vascular tone (ACE). Each graph includes five sampling phases: Pre-Launch (baseline), Post-Launch Suborbital, and ISS Days 4, 7, and 10. For all genes shown, Kruskal–Wallis tests yielded non-significant p-values (p > 0.05), indicating no statistically meaningful expression changes across microgravity exposure. Variability between individuals is represented by standard deviation error bars. These data suggest that certain pathways—including telomerase expression, select cytokines, and components of angiogenic or metabolic signaling—remain transcriptionally stable under the spaceflight conditions studied (p < 0.05, Kruskal-Wallis).

In contrast to the widespread suppression observed in core longevity, mitochondrial, and DNA-repair pathways, the genes represented in this heatmap exhibited minimal or inconsistent transcriptional responses to microgravity. Key metabolic and inflammatory transcripts—including APOE, IL-6, IL-7, and FTO—remained largely unchanged across all flight phases, reflecting the relative stability of systemic metabolic and cytokine networks in circulating immune cells during spaceflight (**Fig. 8**). Genes involved in angiogenesis (VEGFC, VEGFD), vascular signaling (ACE), and growth factor pathways (FGF2, EGF) showed similarly stable expression, suggesting that microgravity does not robustly perturb transcriptional programs governing extracellular remodeling or vascular growth within peripheral leukocytes. Among telomere-related genes, TERT remained consistently low, whereas LMNA displayed moderate fluctuations with a transient increase at ISS Day 7, followed by partial normalization. This pattern, although not statistically significant, may reflect dynamic adjustments in nuclear architecture or mechanosensitive lamin regulation under microgravity. Cell-cycle and ECM-associated transcripts—CDKN1A, TIMP1, and SERPINE1—showed variable but modest changes across time points, while RAP1A (a Ras-family GTPase linked to cell adhesion and immune signaling) exhibited a distinct spike immediately after suborbital launch, potentially reflecting early mechanotransduction or stress-signaling activation before returning toward baseline during ISS sampling. Collectively, these findings indicate that while microgravity profoundly alters genes involved in DNA repair, oxidative stress, mitochondrial function, and senescence, a subset of metabolic, telomere-associated, angiogenic, and ECM-modulating genes is comparatively resilient, showing either stable or non-directional expression patterns. This selective stability suggests pathway-specific thresholds for gravitational sensitivity and highlights the compartmentalized nature of the transcriptional response to spaceflight.

**Fig. 8.**
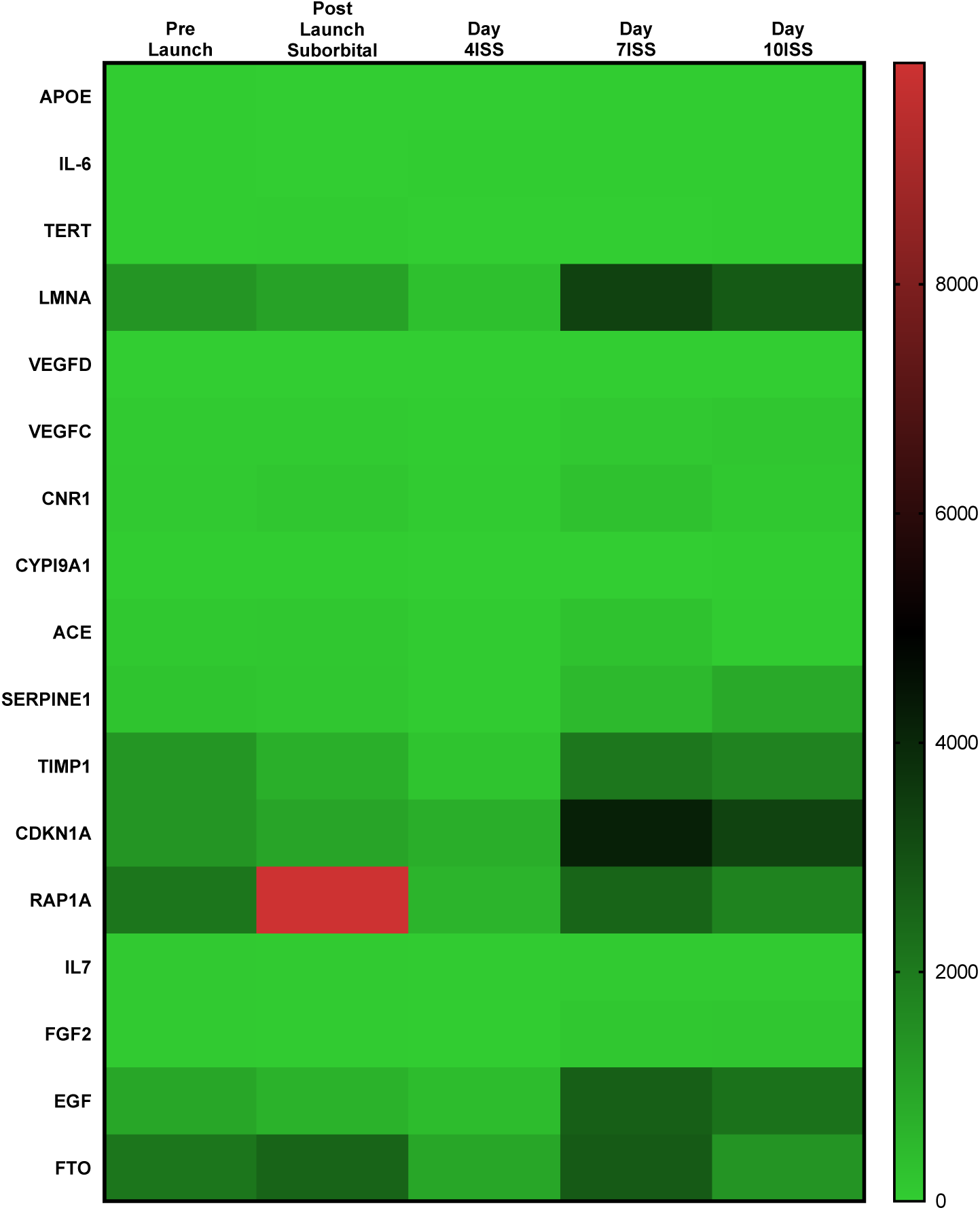
Heatmap of metabolic, inflammatory, angiogenic, telomere-related, and extracellular matrix genes showing minimal or heterogeneous transcriptional responses across microgravity conditions. This heatmap displays normalized RNA-Seq expression values for 17 genes spanning metabolic regulation (APOE, FTO), inflammatory cytokines (IL-6, IL-7), telomere biology (TERT, LMNA), angiogenic/growth factors (VEGFD, VEGFC, EGF, FGF2), xenobiotic metabolism (CYP1A1), vascular signaling (ACE), extracellular matrix remodeling (SERPINE1, TIMP1), and cell-cycle regulation (CDKN1A, RAP1A). Expression levels are shown across five mission phases: Pre-Launch (baseline), Post-Launch Suborbital, and ISS Days 4, 7, and 10. The predominantly green shading indicates low to moderate expression with limited fold changes across microgravity exposure. Notable exceptions include a transient increase in RAP1A expression after suborbital flight and dynamic fluctuations in LMNA, CDKN1A, and TIMP1, although these changes do not follow a consistent directional trend. Overall, the heatmap illustrates that most genes in these pathways remain transcriptionally stable during both short-and intermediate-duration microgravity exposure.

STRING network analysis of genes with non-significant transcriptional changes provides further insight into the selective nature of microgravity-driven molecular remodeling **(Supplementary Fig. 1)**. Although genes such as APOE, FTO, CNR1, and ACE occupy central nodes within metabolic and vascular signaling networks, their expression remained unchanged across all flight phases, suggesting that these systems maintain transcriptional stability despite gravitational unloading. Inflammatory cytokines IL6 and IL7, along with oxidative stress regulator SOD2, also displayed stable expression patterns, consistent with the notion that systemic inflammatory activation is not transcriptionally triggered in circulating leukocytes during the microgravity exposure studied. Similarly, angiogenesis-related genes, including VEGFC, VEGFD, EGF, and FGF2, showed robust interaction networks but no directional expression shifts, indicating that gravitational stress does not broadly influence vascular growth signaling at the transcriptional level. Nuclear architecture and cell-cycle regulators LMNA and CDKN1A, as well as extracellular matrix genes SERPINE1 and TIMP1, exhibited prominently interconnected interaction networks but remained transcriptionally stable, further supporting the idea that microgravity selectively targets specific molecular axes—such as mitochondrial regulation, longevity pathways, and DNA repair—rather than inducing global transcriptional disruption. Finally, telomere-associated TERT and signaling mediator RAP1A formed interaction networks linked to genome stability and immune communication, yet displayed minimal transcriptional fluctuations. This aligns with previous observations that TERT expression often remains epigenetically constrained in peripheral immune cells and may not respond acutely to short- or mid-duration microgravity. Overall, this network-level analysis shows that although these genes occupy biologically important positions within their respective pathways, their transcriptional resilience highlights microgravity’s pathway-specific effects, reinforcing the conclusion that only certain molecular systems—primarily those related to longevity, DNA repair, mitochondrial biology, and immune stress undergo significant reprogramming during spaceflight.

### Telomere Length Changes in Microgravity

Telomere length was quantified in peripheral blood samples collected pre-launch and during ISS Days 4 and 10 using the Absolute Human Telomere Length Quantification qPCR Assay Kit and analyzed by the comparative ΔΔCq approach (**Fig. 10**). Compared with baseline, telomere length displayed a significant elongation on ISS Day 4 (p < 0.05), indicating an acute early-flight response to microgravity. By Day 10, telomere length exhibited a decreasing trend relative to Day 4; however, values remained above pre-launch levels (p = 0.07), suggesting a partial reversion toward baseline while maintaining an overall elongation state. This biphasic pattern—initial elongation followed by gradual decline—aligns with reports from the NASA Twins Study^8^ in which telomere lengthening occurred during spaceflight and was subsequently followed by post-flight shortening. The early elongation observed here may reflect microgravity-driven alterations in oxidative stress burden, enhanced DNA repair activity, or activation of telomerase-mediated protective mechanisms^8,9^. The subsequent attenuation by Day 10 suggests the emergence of compensatory homeostatic processes as cells adapt to the microgravity environment. Overall, these findings indicate that short-term microgravity exposure induces a significant but transient elongation of telomeric regions, highlighting the sensitivity of telomere biology to acute spaceflight-associated physiological changes.

**Fig. 10.**
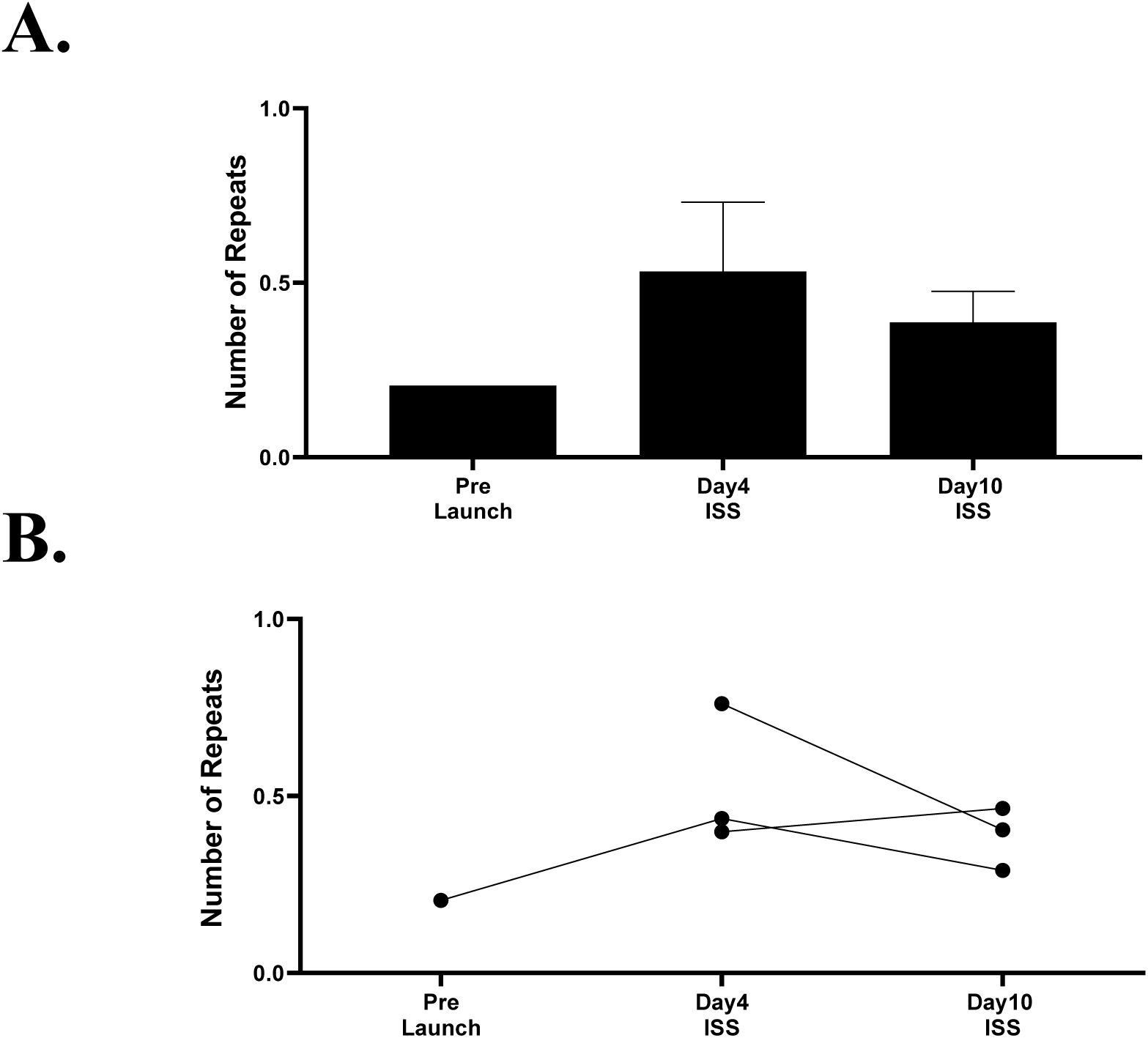
Telomere length dynamics during microgravity exposure. **(A)** Group-level changes in telomere length measured in peripheral blood across three time points: Pre-Launch (n=1), ISS Day 4 (n=3), and ISS Day 10 (n=3). Telomere length was quantified using the Absolute Human Telomere Length Quantification qPCR Assay Kit and calculated by the comparative ΔΔCq method. Bars represent mean values with standard deviation. **(B)** Individual astronaut trajectories illustrating within-subject telomere length variation across flight phases. A significant increase in telomere length was observed on ISS Day 4 compared with pre-launch values (p < 0.05). By ISS Day 10, telomere length declined relative to Day 4 but remained elevated above baseline (p = 0.07). These results demonstrate a transient elongation response during early microgravity exposure, followed by partial normalization over time.

### Microgravity induces strong suppression of miRNAs linked to cell cycle control and DNA repair

Analysis of microRNA expression revealed a robust decrease in several miRNAs known to regulate proliferation, senescence, and DNA damage pathways. MiR-17HG—representing the miR-17-92 cluster host gene—showed a significant decline immediately after suborbital ascent and remained markedly reduced throughout ISS exposure (p = 0.0334) (**Fig. 11**). A similar pattern was observed for miR-29B2CHG (p = 0.0233) and miR29A (p = 0.0016), both of which exhibited strong downregulation by Day 4 and remained suppressed on Days 7 and 10. Notably, these miRNAs play critical roles in modulating DNA repair enzymes (e.g., PARP1, ATM), collagen remodeling, mitochondrial metabolism, and apoptosis signaling, suggesting broad functional consequences of their microgravity-responsive repression^10,13^. In contrast, miR-34A and miR-21 were expressed at very low levels across all conditions, with no significant fluctuations during flight. The heatmap visualization highlights the coordinated suppression of the miR-17-92 and miR-29 families specifically during in-flight ISS timepoints, supporting the concept of a microgravity-driven shift toward reduced proliferative signaling and altered stress-response regulation. Collectively, these findings indicate that short-duration spaceflight markedly attenuates key miRNA networks involved in maintaining genome integrity and cellular adaptation, aligning with the observed downregulation of DNA repair genes and modulation of telomere dynamics in the same astronaut samples^8,9^.

**Figure 11.**
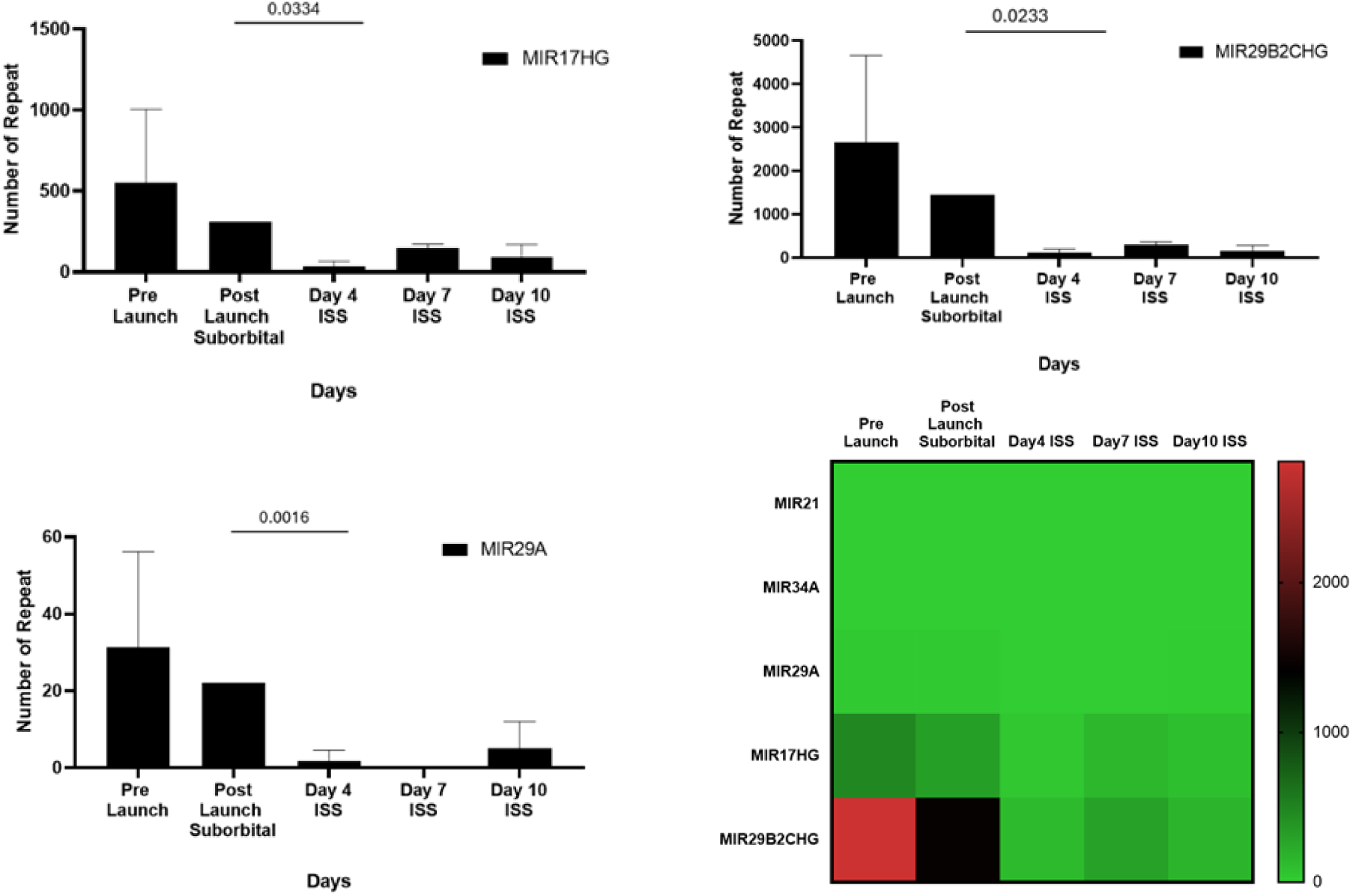
Microgravity-Associated Downregulation of miRNA Clusters Involved in Cell Cycle, Senescence, and DNA Repair Pathways. Expression levels of five microgravity-responsive microRNAs (miR-17HG, miR-29B2CHG, miR-29A, miR-34A, and miR-21) were quantified across five mission timepoints: Pre-Launch, Post-Launch Suborbital, Day 4 ISS, Day 7 ISS, and Day 10 ISS. Bar plots display mean copy numbers ± SD, and significance values are indicated above relevant comparisons. MiR-17HG (p = 0.0334), miR-29B2CHG (p = 0.0233), and miR-29A (p = 0.0016) showed significant reductions relative to Pre-Launch. miR-34A and miR-21 exhibited consistently low abundance across all timepoints. The accompanying heatmap summarizes overall patterns of miRNA suppression during ISS exposure, demonstrating a pronounced decrease in members of the miR-17-92 and miR-29 families—key regulators of proliferation, DNA damage responses, mitochondrial homeostasis, and ECM remodeling.

## 4. Discussion

Microgravity, as a unique physical stressor encountered in space, alters fundamental biological processes, including cell proliferation, DNA damage response, mitochondrial function, immune signaling, and telomere dynamics^8,9,11^. Aging biology, which is inherently sensitive to environmental perturbations, has been shown to respond dramatically under spaceflight conditions^1,8^. In this study, we investigated how microgravity influences the transcriptional and functional landscape of longevity-related genes in human immune cells, providing the first multi-layered evidence based on Turkiye’s first manned space mission. One of the most striking findings of this study was the dynamic alteration in telomere length. A transient elongation observed on Day 4 was followed by a decrease toward baseline levels by Day 10, echoing the findings of the NASA Twins Study, where telomere elongation during spaceflight and post-flight erosion were similarly reported ^8^. These results support the model that telomere biology is not only a function of cellular aging but is acutely sensitive to micro-environmental stress, such as microgravity and radiation exposure.

The transient telomere elongation observed during early ISS exposure appears to occur within a broader landscape of microgravity-induced molecular reprogramming that includes suppression of AP2A1-centered endocytic signaling, widespread downregulation of DNA repair genes, and shifts in miRNA regulatory networks. Early telomere lengthening may represent a compensatory response to reduced mechanotransduction stress, consistent with our finding that AP2A1 and associated clathrin–integrin trafficking components are markedly downregulated in microgravity. Given that integrin signaling and cytoskeletal tension influence telomere positioning, nuclear architecture, and shelterin function, the suppression of AP2A1-mediated mechanotransduction may transiently relax the biophysical constraints on chromosome ends, allowing elongation despite reduced repair capacity^17,21^. However, the simultaneous downregulation of critical DNA repair genes such as WRN, BRCA1/2, ATM, and TP53 suggests that telomere homeostasis becomes increasingly vulnerable as exposure persists, potentially explaining the decline in telomere length observed by Day 10. Microgravity-responsive miRNAs—particularly miR-34a, miR-21, and the miR-30 family, all of which target SIRT1, MTOR, and ATM—likely amplify these effects by coordinately suppressing nutrient-sensing, mitochondrial, and genome maintenance pathways known to stabilize telomeres^9,10,14^. Together, these findings support a model in which early microgravity reduces mechanical stress and triggers transient telomere elongation, but progressive suppression of DNA repair and mechanotransduction pathways, reinforced by miRNA-mediated repression, drives a return toward baseline telomere length as cellular homeostasis attempts to re-establish equilibrium. This integrated framework highlights telomeres as a sensitive readout of the combined mechanical, metabolic, and genomic stresses imposed by spaceflight. Transcriptomic analysis revealed significant downregulation of several core longevity-associated genes, including SIRT1, MTOR, TP53, WRN, and NFE2L2. The downregulation of SIRT1, a key NAD-dependent deacetylase regulating DNA repair and mitochondrial function, may compromise genome stability and cellular homeostasis in microgravity ^22^. Interestingly, the suppression of MTOR, a master regulator of nutrient signaling and cell growth, aligns with studies suggesting that mTOR inhibition under space stress acts as a protective metabolic adaptation ^23^. The decrease in TP53 and WRN—both pivotal in genome integrity—further underscores microgravity’s effect on DNA repair mechanisms. Importantly, antioxidant genes SOD2 and NFE2L2 were downregulated, potentially impairing the cell’s ability to manage reactive oxygen species (ROS), which are elevated in spaceflight environments ^19,24^.

Many of these genes are regulated through long-term endocrine, metabolic, or tissue-specific control mechanisms rather than rapid stress-response circuits, making them less responsive to short-duration microgravity exposure in circulating immune cells^11,18^. For example, TERT expression in leukocytes is typically low and tightly epigenetically constrained, meaning that telomerase activation reflects chronic proliferative or inflammatory stimuli rather than transient environmental stressors^1,8^. Likewise, angiogenic genes such as VEGFC/D and EGF are predominantly regulated by tissue-level hypoxia and stromal signaling, conditions not directly mirrored in peripheral blood transcriptional profiles during spaceflight^8,11^. The unchanged cytokines IL-6 and IL-7 may reflect the complexity of immune regulation in vivo, where circulating transcript levels do not necessarily track with protein secretion or localized lymphoid activation. Genes such as APOE, ACE, and FTO are largely governed by metabolic, hepatic, or neuroendocrine regulatory networks, which are slower to exhibit transcriptional drift and may require longer-duration missions or chronic unloading to manifest measurable changes^8,11^. These stable expression patterns highlight that microgravity exerts pathway-selective pressure, profoundly affecting mechanotransduction, DNA repair, mitochondrial quality control, and inflammatory stress pathways while leaving other metabolic or endocrine networks transcriptionally resilient. This differential sensitivity underscores the compartmentalized and hierarchical nature of microgravity adaptation, in which only certain biological systems undergo rapid transcriptomic remodeling, whereas others maintain homeostatic stability despite environmental unloading. The coordinated downregulation of AP2A1 and other endocytic machinery components under microgravity has important implications for cellular aging, structural integrity, and mechanotransduction^17,21^.

Clathrin-mediated endocytosis is tightly coupled to actin dynamics and membrane tension sensing; thus, suppression of AP2 complex genes may reflect a broader collapse of cytoskeletal–membrane communication networks in gravitational unloading. Recent studies have demonstrated that AP2A1 serves as a mechanosensitive regulator capable of shifting cell states between senescence and rejuvenation, in part through its interactions with integrin β1 along actin stress fibers ^21^. Reduced AP2A1 expression—similar to that observed in our astronauts—has been shown to preserve cytoskeletal resilience and limit senescence-associated signaling, suggesting that microgravity may transiently promote a rejuvenation-like transcriptional state by dampening integrin–cytoskeleton feedback loops^17,21^. Furthermore, impaired clathrin–actin coupling can disrupt nutrient sensing, mitochondrial dynamics, and inflammatory signaling, all of which are established hallmarks of aging^1,17^. As mechanotransduction pathways such as focal adhesion kinase (FAK), mTOR, and YAP/TAZ are gravity-sensitive, the suppression of adaptor proteins involved in endocytosis may represent an upstream trigger for the altered metabolic, oxidative, and genomic stability programs observed in spaceflight^11,17^. Therefore, the downregulation of AP2-family genes under microgravity may not simply reflect reduced vesicle trafficking but a fundamental recalibration of structural and signaling networks that anchor cellular aging processes to mechanical forces.

The suppressed expression of PINK1 and UCP1, involved in mitochondrial maintenance, is also consistent with findings that report compromised mitochondrial function across tissues in space ^8^. This multi-gene suppression paints a systemic picture of oxidative stress and mitochondrial dysfunction as major hallmarks of space-induced cellular aging. In contrast, IL15 was upregulated, suggesting an adaptive immune response potentially maintaining NK and homeostatic T-cell function, while IL6, IL1B, and IFNG were downregulated, indicating a general trend toward immune suppression, in agreement with previous microgravity studies ^20^. The coordinated molecular patterns observed in this study suggest that telomere elongation, AP2A1 stability, DNA repair signaling, and microgravity-responsive miRNAs are mechanistically intertwined components of the astronaut adaptive response. The early elongation of telomeres on Day 4 of ISS exposure aligns with prior evidence that microgravity transiently reduces oxidative stress and activates compensatory DNA repair and telomerase-associated pathways, producing a temporary genomic “protective” phenotype^8,9^. Consistent with this, AP2A1—an adaptor protein linked to endocytic trafficking, cellular stress buffering, and chromatin accessibility^21,22^—remained strikingly stable throughout spaceflight, indicating preservation of vesicular transport and intracellular signaling during gravitational unloading. This stability may protect against excessive DNA damage, thereby supporting the observed telomere maintenance response. However, the significant suppression of miRNAs belonging to the miR-17-92 and miR-29 families suggests a parallel dampening of pathways controlling cell cycle progression, apoptosis, and DNA repair enzyme expression. Notably, miR-29 family members normally regulate key DNA repair genes such as PARP1 and ATM, while miR-17HG-driven clusters control proliferation and genomic stability. Their reduction during flight may reflect an adaptive shift toward minimizing proliferative signaling under mechanical unloading, thereby reducing replication-associated telomere attrition and conserving cellular resources in microgravity. Together, these findings imply that microgravity induces a unique molecular state characterized by telomere extension, maintained AP2A1-mediated signaling homeostasis, and suppression of miRNAs that ordinarily drive DNA-repair and growth-associated programs. This integrated response may represent a finely tuned balance between protective adaptation and controlled downregulation of energy-intensive genomic maintenance pathways, offering new insights into how human cells modulate longevity, genomic stability, and mechanotransductive signaling during spaceflight. Beyond statistically significant genes, several others displayed biologically meaningful expression shifts over time. For example, TERT, FGF2, and VEGFC exhibited dynamic, non-monotonic expression patterns. Despite not crossing the conventional p-value threshold, these patterns may indicate early-stage or transient regulatory responses. This observation supports the growing consensus that p-values alone are insufficient to capture biological relevance, especially in complex systems exposed to multifactorial stressors like microgravity. STRING-based network analysis further illustrated that downregulated genes—such as SIRT1, TP53, and NFE2L2—are tightly connected within longevity and stress resistance modules. These network patterns reinforce the concept that spaceflight disrupts coordinated gene programs rather than isolated transcripts. Such systems biology perspectives are vital, as highlighted by Zhao et al. (2023), who emphasized the importance of integrating interaction networks to interpret transcriptomic adaptations under stress conditions ^18,25^. The transcriptional stability of APOE, IL-6, TERT, VEGFC/D, FTO, ACE, and other unchanged genes suggests that not all physiological pathways exhibit gravity-dependent sensitivity, and that certain molecular systems may be buffered against acute environmental perturbation.

The broad downregulation of longevity-associated transcripts observed in the heatmap aligns closely with the AP2A1-centered suppression of endocytic machinery, suggesting that microgravity triggers a unified collapse of cytoskeletal, membrane-trafficking, and genomic stability pathways. Reduced expression of AP2A1 and its clathrin-interacting partners may impair integrin recycling, mechanotransduction, and mitochondrial–cytoskeletal coupling—processes known to regulate telomere maintenance and chromatin architecture^1,17,21^. This is consistent with recent evidence that mechanical unloading weakens telomere-protective pathways, reducing shelterin stability and accelerating telomere attrition through impaired DNA repair signaling^8,9^. The observed downregulation of WRN, BRCA1/2, ATM, and TP53 in our heatmap further supports this model, as these genes are core regulators of telomere replication fidelity and DNA damage sensing. Moreover, several microgravity-responsive miRNAs identified in astronaut studies—including miR-34a, miR-21, miR-30 family members, and miR-424—directly target genes involved in mechanotransduction and telomere maintenance, such as SIRT1, MTOR, and ATM, providing an upstream regulatory layer that may amplify transcriptomic suppression under microgravity. Together, these interconnected findings indicate that gravitational unloading activates an miRNA-driven regulatory program that diminishes endocytic and cytoskeletal integrity, weakens telomere-protective DNA repair networks, and globally suppresses longevity pathways. This coordinated transcriptional reprogramming suggests that microgravity effects on aging biology are not isolated events but reflect a cohesive systems-level response involving mechanical signaling, epigenetic regulation, and genome maintenance machinery. Collectively, these findings provide a comprehensive model of how microgravity modulates cellular aging and immune responses. The convergence of telomere dynamics, mitochondrial dysfunction, oxidative stress, and altered immune signaling highlights the profound systemic reprogramming induced by spaceflight. Importantly, the observed modulation of AP2A1 aligns with emerging terrestrial evidence of its pivotal role in cellular aging, positioning it as a potential biomarker and therapeutic target not only for astronaut health but also for aging-related interventions on Earth^1,21^. Thus, this integrative transcriptomic-telomeric analysis demonstrates that microgravity acts as a multidimensional environmental modulator of human cellular aging and immunity. These insights are not only foundational for space biology but also hold translational promise for developing countermeasures against aging-related diseases. Ultimately, our findings may guide the design of personalized monitoring and intervention strategies to protect human health during long-term space travel while simultaneously informing novel anti-aging strategies on Earth.

## Conclusion

In summary, this study provides integrative evidence that short-duration spaceflight induces a coordinated remodeling of longevity-associated molecular networks in human peripheral blood cells. The convergence of transient telomere elongation, widespread suppression of DNA repair and endocytic mechanotransduction genes, and attenuation of miRNA programs governing genome stability, metabolism, and stress adaptation suggests that microgravity elicits a distinct, time-limited reprogramming of cellular homeostasis rather than uniform molecular damage. Central to this response is the downregulation of AP2A1 and associated clathrin-mediated trafficking components, which likely alters integrin–cytoskeleton signaling, chromatin organization, and mitochondrial–nuclear communication, thereby reshaping aging-relevant pathways under mechanical unloading. The early telomere elongation observed during orbital exposure, followed by partial normalization, supports a model in which microgravity transiently relaxes biophysical and regulatory constraints on chromosome ends, potentially generating a short-lived genomic “protective” state despite concurrent dampening of canonical DNA repair programs. Collectively, these findings position AP2A1-centered mechanotransduction and miRNA-mediated regulation as key modulators of spaceflight-induced aging phenotypes and highlight their broader relevance as biomarkers and therapeutic targets for promoting resilience to mechanical stress, both in astronauts and in age-related disorders on Earth.

## Author Contributions

E.C.: Data Collection, analysis, figure creation, transcriptome interpretation, and manuscript writing. O.D.: Data collection, analysis, and transcriptome interpretation. B.T.: Telomere analysis, longevity gene identification, and figure design. D.B.: miRNA transcriptome analysis and figure design. E.G.: Data visualization and figure design. S.N.H.E.: Telomere analysis and data interpretation. B.Y.: Software support and transcriptomic analysis. K.K.: Bioinformatics analysis and transcriptomic processing. I.H.K.: Laboratory infrastructure and telomere analysis support. F.K.: Transcriptomic analysis interpretation and manuscript writing. B.A.: STRING analysis, Graphic design, data interpretation, and manuscript writing. C.T.: Study supervision, experimental planning and analysis, project coordination, and manuscript writing.

## Acronyms/Abbreviations

MESSAGE: (Microgravity Associated Genetics) Science Mission;
International Space Station: (ISS);
Peripheral Blood Mononuclear Cells: (PBMC);
Scientific and Technological Research Council of Turkiye: (TUBITAK);
Turkish Space Agency: (TUA);
Quantitative PCR: (qPCR);
RNA Sequencing: (RNA-Seq);
Transgenic Cell Technologies and Epigenetic Application and Research Center: (TRGENMER)

## Funding Acknowledgement

This study was supported by the Scientific and Technological Research Council of Turkiye (TUBITAK) and the Turkish Space Agency (TUA) through the Turkish Astronaut and Scientific Mission (TABM) Project (1007-KAMAG-121L002 / Agreement No: TABM-HZT-23-25). We would like to thank TUBITAK UZAY and TUA for their support and declare that none of the opinions and findings contained in this publication are the official views of TUBITAK and TUA.

## Acknowledgements

The experience and infrastructure gained through these efforts provided the foundation for participation in the MESSAGE Science Mission. We gratefully acknowledge the support of TUA, TUBITAK, and Uskudar University throughout this process. The opinions and findings expressed in this publication are solely those of the authors and do not necessarily represent the official views of the supporting institutions. Finally, we extend our sincere thanks to all research colleagues, data analysts, and technical staff whose contributions were essential to the success of the MESSAGE Science Mission.

## Data Availability

The datasets generated and analyzed during the current study are available from the corresponding author upon reasonable request.

## Supplementary Figures

**Supplementary Figure 1.**
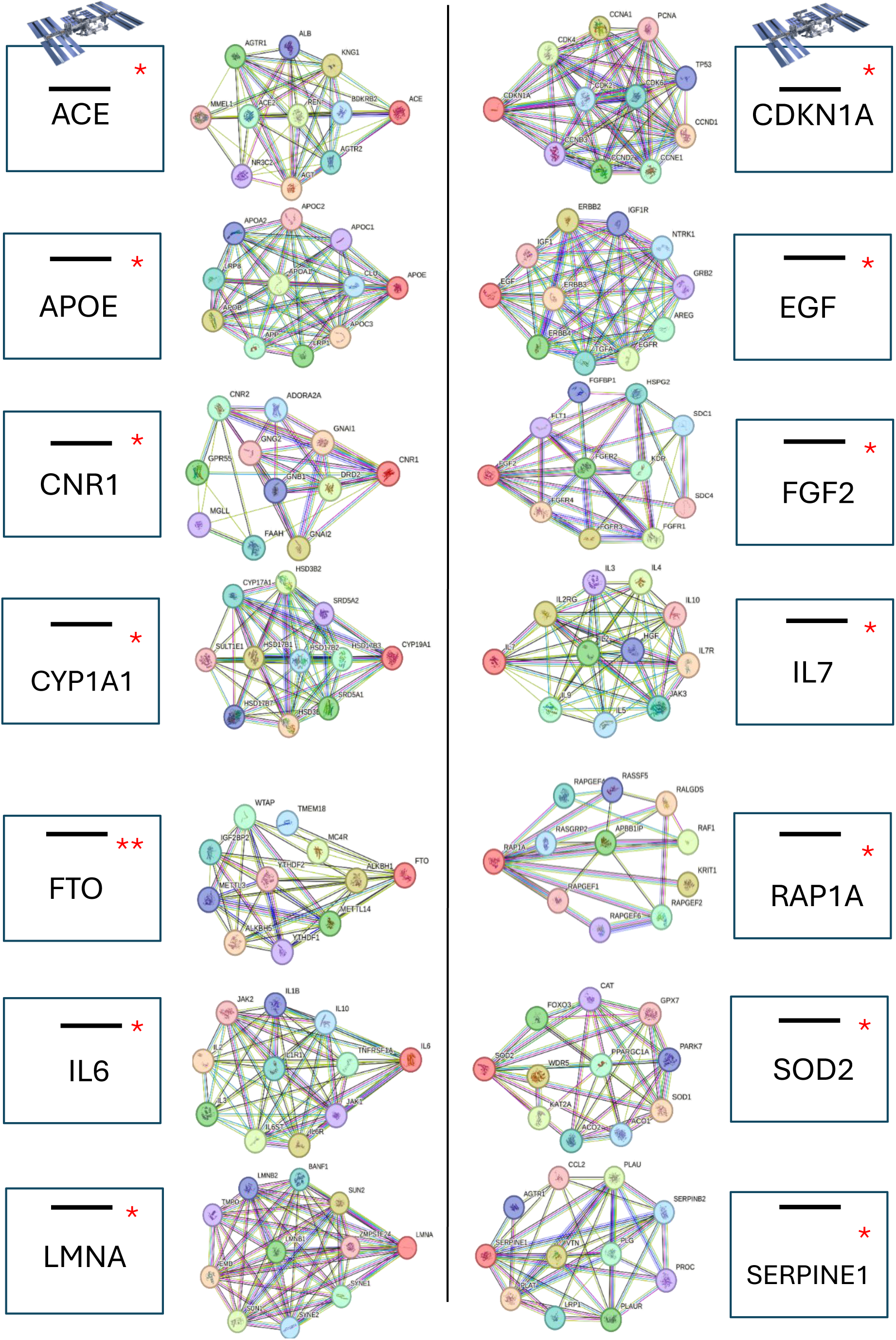
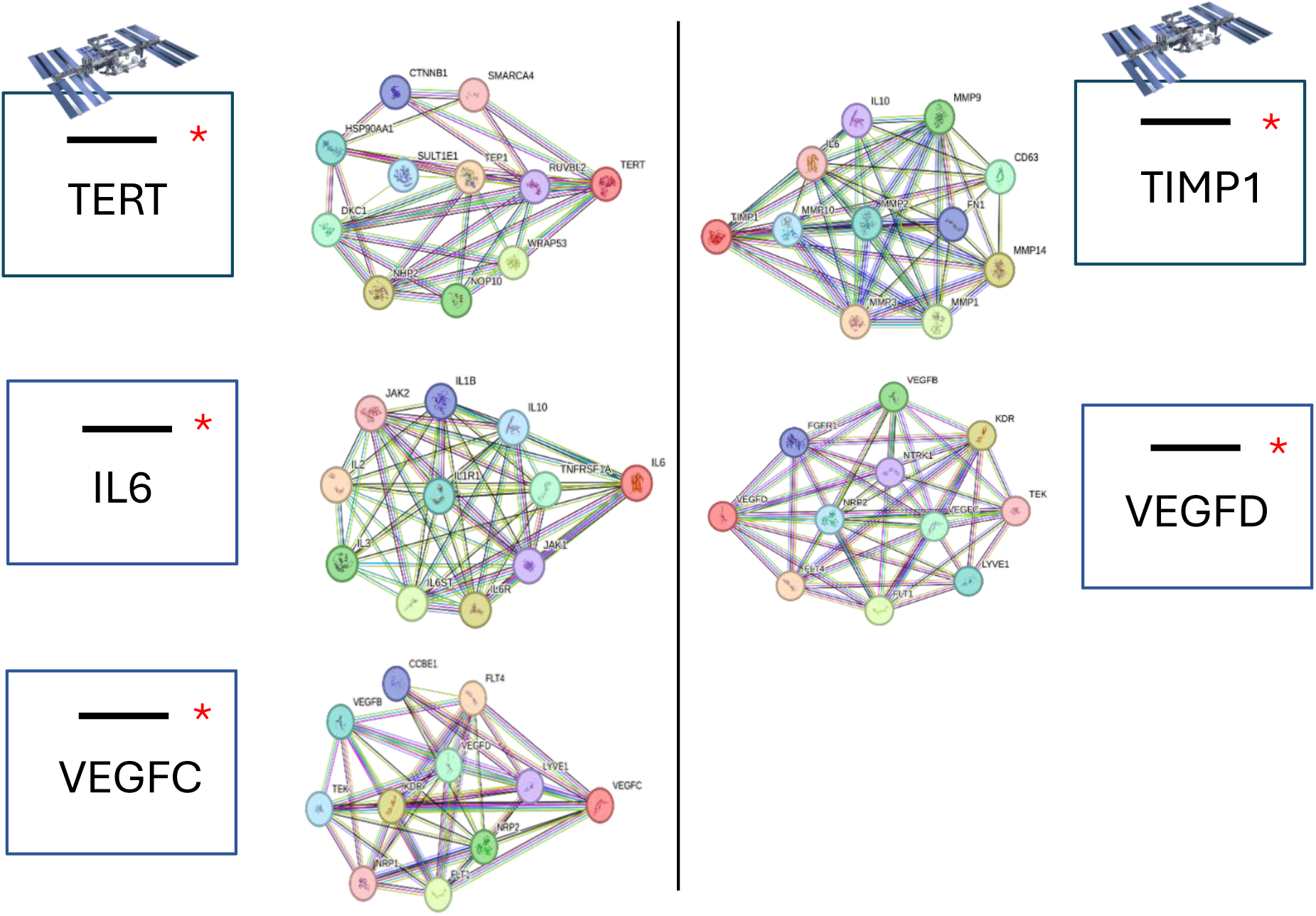
STRING protein–protein interaction (PPI) networks for genes exhibiting minimal or non-significant transcriptional changes across suborbital and orbital microgravity exposure. This figure shows STRING-generated interaction networks for metabolic, inflammatory, angiogenic, telomere-associated, and extracellular matrix–regulating genes that did not demonstrate statistically significant expression changes in RNA-Seq analysis. Each panel highlights the interaction partners for genes involved in: Metabolism and neuroendocrine signaling: APOE, FTO, CNR1; Inflammation and cytokine regulation: IL6, IL7; Angiogenesis and vascular signaling: VEGFC, VEGFD, EGF, FGF2, ACE; Cell-cycle and nuclear architecture: CDKN1A, LMNA; Oxidative stress resilience and detoxification: CYP1A1, SOD2; Extracellular matrix remodeling: SERPINE1, TIMP1; Telomere maintenance pathways: TERT; GTPase-mediated signaling: RAP1A. Black lines indicate no significant change in gene expression. The dense networks illustrate known or predicted functional associations, including co-expression, experimental evidence, and curated pathway interactions. Despite these genes’ involvement in broad signaling modules, their transcriptional stability suggests that microgravity-induced changes primarily target other molecular systems rather than these pathways.

